# A Single-Short Partial Reprogramming of the Endothelial Cells decreases Blood Pressure via attenuation of EndMT in Hypertensive Mice

**DOI:** 10.1101/2024.05.20.595057

**Authors:** Laena Pernomian, Emily W. Waigi, Vi Nguyen, Ahmed D. Mohammed, Tiago J. da Costa, Milene T. Fontes, Jason L. Kubinak, Andrew Aitken, Vinicia Campana Biancardi, David A. Sinclair, Cameron G. McCarthy, Yunguan Wang, Wenbin Tan, Camilla Ferreira Wenceslau

## Abstract

**Background:** Small artery remodeling and endothelial dysfunction are hallmarks of hypertension. Growing evidence supports a likely causal association between cardiovascular diseases and the presence of endothelial-to-mesenchymal transition (EndMT), a cellular transdifferentiation process in which endothelial cells (ECs) partially lose their identity and acquire additional mesenchymal phenotypes. EC reprogramming represents an innovative strategy in regenerative medicine to prevent deleterious effects induced by cardiovascular diseases.

**Methods:** Using a partial reprogramming of ECs, via overexpression of Oct-3/4, Sox-2, and Klf-4 (OSK) transcription factors, we aimed to bring ECs back to a youthful phenotype in hypertensive mice. Primary ECs were infected with lentiviral vectors (LV) containing the specific EC marker cadherin 5 (Cdh5) and the fluorescent reporter enhanced green fluorescence protein (EGFP) with empty vector (LVCO) or with OSK (LV-OSK). Confocal microscopy and western blotting analysis were used to confirm the OSK overexpression. Cellular migration, senescence, and apoptosis were evaluated. Human aortic ECs (HAoECs) from male and female normotensive and hypertensive patients were analyzed after OSK or control treatments for their endothelial nitric oxide synthase (eNOS) levels, nitric oxide (NO), and genetic profile. Male and female normotensive (BPN/3J) and hypertensive (BPH/2J) mice were treated with an intravenous (i.v.) injection of LVCO or LV-OSK and evaluated 10 days post-infection. The blood pressure, cardiac function, vascular reactivity of small arteries, *in vivo* EGFP signal and EndMT inhibition were analyzed.

**Results:** OSK overexpression induced partial EC reprogramming *in vitro*, and these cells showed endothelial progenitor cell (EPC)-like features with lower migratory capability. OSK treatment of hypertensive BPH/2J mice normalized blood pressure and resistance arteries hypercontractility, via the attenuation of EndMT and elastin breaks. EGFP signal was detected *in vivo* in the prefrontal cortex of both BPN/3J and BPH/2J-treated mice, but OSK induced angiogenesis only in male BPN/3J mice. OSK-treated human ECs from hypertensive patients showed high eNOS activation and NO production, with low ROS formation. Single-cell RNA analysis showed that OSK alleviated EC senescence and EndMT, restoring their phenotypes in human ECs from hypertensive patients.

**Conclusion:** Overall, these data indicate that OSK treatment and EC reprogramming can decrease blood pressure and reverse hypertension-induced vascular damage.

## Introduction

The major pathophysiological characteristic of hypertension is the occurrence of small artery remodeling and endothelial dysfunction [1,2]. Until recently, endothelial dysfunction had been primarily considered as imbalanced vasodilation and vasoconstriction, characterized by a reduction in nitric oxide (NO) bioavailability, elevated reactive oxygen species (ROS), and exacerbated production of pro-inflammatory mediators. However, growing evidence supports a robust and likely causal association between cardiovascular diseases and the presence of endothelial-to-mesenchymal transition (EndMT) [3,4,5,6,7], a cellular transdifferentiation process in which endothelial cells (ECs) partially lose their identity and acquire additional mesenchymal phenotypes, including the gain of contractile properties.

Under physiological conditions, ECs display remarkable phenotypic plasticity [8,9,10]. They can express features of alternative EC lineages, with the phenotypes being regulated by genetic and environmental stimuli. However, adverse conditions such as cardiovascular diseases can trigger the formation of EndMT-induced ECs which do not transdifferentiate back to their original phenotype with a normal function, and in chronic stages, they lead to perturbations in the vascular function. The lack of information in the literature about EndMT in essential hypertension makes it difficult to understand whether this phenotypic transitioning of ECs contributes to hypertension-induced damage in the cardiovascular system. We hypothesized that arteries from hypertensive mice present high levels of EndMT and that specific EC reprogramming can decrease blood pressure values and restore vascular dysfunction in resistance arteries in hypertensive mice. Here, we have observed that EndMT is exacerbated in hypertension. There were increased internal elastic lamina (IEL) breaks in resistance arteries from spontaneously hypertensive mice, indicating a potential migration of EndMT-induced ECs to the media layer. This was evidenced by the presence of EndMT in the intima and media layers, which can further contribute to the deleterious vascular damage observed in hypertension. By taking advantage of this extraordinary plasticity, we then manipulated ECs’ fate by inducing a single-short partial cellular reprogramming, which consisted of only one exposure to specific transcription factors, without the ECs passing through the pluripotent state. For this, we overexpressed three master transcriptional factors Oct-3/4, Sox-2, and Klf-4 (OSK) [11,12,13] in ECs. Reprogrammed ECs *in vitro* exhibited endothelial progenitor cell (EPC)-like features, with low migration capability and reduced cellular senescence phenotype that prevented EndMT formation. For human ECs, the forced expression of OSK induced NO synthesis and decreased ROS generation in male and female human ECs from hypertensive patients. Single-cell RNA analysis showed that OSK alleviated EC senescence and EndMT, facilitating the restoration of EC phenotypes in human hypertensive ECs. Reprogrammed ECs *in vivo* restored blood pressure in male hypertensive mice, and improved vascular contractility and remodeling by preventing elastin breaks.

## Methods

### Lentiviral vector (LV) production

For more information, please refer to Supplemental Methods. Human embryonic kidney (HEK293T) cells were cultured and transfected with three packaging plasmids (psPAX2, pRSV-Rev, and pMD2.G), and the lentiviral vector (LV) for mouse Cadherin 5-(also known as Cdh5 or VE-cadherin)-Oct3/4-Sox2-Klf4-EGFP (here, referred as LV-OSK) or the Cdh5-EGFP (referred as LVCO). LV production and utilization in cells and mice had the School of Medicine University of South Carolina ethics committee approval (IACUC #2596-101690-041122, and IBC protocol # 300322).

### Mouse and Human EC culture

C57Bl/6 mouse intestinal mesenteric primary endothelial cells (Accegen, ABC-TC3197) were used in the following experiments with phosphate-buffered saline (PBS), LVCO or LV-OSK, and evaluated 3- or 5-days post-treatments.

The primary normotensive and hypertensive human aortic endothelial cell (HAoEC) were purchased from PromoCell, and infected with sendai virus (SeV) carried polycistronic Klf4-Oct3/4-Sox2 and Klf4 vectors (ThermoFisher, Waltham, MA, #A16517), here referred as OSK-SeV, to induce cell reprogram, or with control SeV with EmGFP (Thermo Fisher Scientific, A16519), named as EGFP-SeV (control condition), and the HAoECs were collected at day 5 for further analysis.

### Animals

All animal procedures and protocols used were approved by the Animal Care and Use Committee at the University of South Carolina School of Medicine. Experiments were conducted following the National Institutes of Health Guide for the Care and Use of Laboratory Animals and Animal Research Reporting of in Vivo Experiments (ARRIVE) guidelines. Male and female BPN/3J (RRID:IMSR_JAX:003004) and BPH/2J (RRID:IMSR_JAX:003005) mouse strains were obtained from The Jackson Laboratory and maintained as an inbred colony at the Animal Facility (School of Medicine Columbia, University of South Carolina). Male and female mice were used at 40-44 weeks of age for the intravenous treatment, and a female mice group at 30 weeks of age was used for the intraperitoneal treatment with the LVCO or LV-OSK. All mice were maintained on a 12-hour light/dark cycle with water and standard chow diet ad libitum.

### Flow cytometry

Treated mouse endothelial cells were analyzed by flow cytometry for their EGFP positive (EGFP^+^) signals, or for activation of apoptosis and necrosis, using the Pacific Blue Annexin V Apoptosis Detection Kit with 7-amino-actinomycin D (7-AAD) (BioLegend, #640926), according to the manufacturer’s instructions.

### Wound healing assay

PBS-, LVCO-, or LV-OSK-treated mouse endothelial cells (3 days) were plated on a 24-well plate with a fresh endothelial cell medium, and the wound healing assay was performed by scratching on the bottom of the well plate. Photos were taken after scratch and were referred as 0 h, and after 24 h to check for cellular growth.

### Confocal microscopy and immunofluorescence

Mouse ECs were treated with PBS, LVCO, or LV-OSK for 3 days (for CD31, CD133, CD34, and CD45 evaluation) or 5 days (for CD31, Oct-3/4, Ki67, Sox-2, and total histone H2Ax) prior to the immunofluorescence protocol. HAoEC were treated with PBS (control) or OSK-SeV (with no EGFP) for 5 days. The coronary arteries in the left ventricle from male BPN/3J or BPH/2J mice with no treatments (at 72 weeks of age), and MRA isolated from male and female BPN/3J or BPH/2J mice also with no treatments (at 72 weeks of age) were used to show the EndMT (CD31, α-SMA and DAPI staining). Brains (prefrontal cortex) isolated from i.v. treated LVCO or LV-OSK male and female mice at 40-44 weeks of age were used to detect *in vivo* EGFP signals (from the LV) and CD31 expression, and thoracic aortas from male mice treated with LVCO or LV-OSK (40-44 weeks of age) were evaluated for CD31, α-SMA; DAPI and EGFP detection by immunofluorescence.

### Western blotting

Mouse endothelial cells treated with PBS, LVCO, or LV-OSK for 3 days were evaluated by Western blotting. Additionally, 5 days post-infection EGFP-SeV (control) or EGFP-OSK-SeV, the normotensive and hypertensive HAoEC were serum-starved for 2 h before the experiment was performed. The cells were analyzed to check their protein expression of Klf-4, phosphorylated Serine 1177 (S1177) of endothelial nitric oxide synthase (eNOS), total eNOS and glutharaldehyde-phosphate dehydrogenase (GAPDH).

### Senescence-associated β-galactosidase assay

Cellular senescence was evaluated in mouse ECs (passage 7) by the Senescence-Associated (SA) β-galactosidase activity assay (Abcam, ab65351) after treatment with PBS, LVCO, or LV-OSK, and processed as manufacturer’s instructions.

### Digital spatial profiling (DSP) of aortas from LVCO and LV-OSK-treated mice at a transcript-proteomic scale

DSP was performed on formalin-fixed paraffin-embedded (FFPE) thoracic aortas isolated from the i.v. LVCO- or LV-OSK-treated (40-44-week-old) mice, and the Mouse Whole Transcriptome Atlas (RNA v1.0) was applied to the samples. The utilization of multicolored morphology markers targeting pan-cytokeratin (Pan-CK; 2 μg/mL; Novus Biologicals; NBP2-33200) to stain filamentous proteins of epithelial cells, CD45 (1:40 dilution; Novus Biologicals; NBP1-44763AF594) as a transmembrane protein of all differentiated hematopoietic cell marker, and SYTO83 (0.2 μM; ThermoFisher Scientific; S11364) for the nucleus, enabled the visualization of different compartments. This visualization guided the selection of regions of interest (ROIs; i.e., the endothelial or the vascular smooth muscle layers). The samples underwent 20x high-precision scanning using a GeoMx DSP system, followed by the selection of ROIs. The DNA oligonucleotides, which were attached to the profiling reagents, were released sequentially using ultraviolet illumination and gathered into individual wells on a 96-well plate. Subsequently, the collected DNA was subjected to Illumina library preparation. The expression levels were measured using an Illumina Sequencer and then analyzed using the DSP interactive software (GeomxTools, version 1.99.4; Advanced Genomics Core).

### RNA sequencing for human and mouse ECs

Mouse ECs were treated for 24 h with TNF-α (100 nM) or vehicle (0.1 M phosphate buffer) or HAoECs from male and female normotensive and hypertensive patients treated with OSK or control SeV, and their genetic profile were analyzed by RNA sequencing.

### Nitric oxide (NO) and reactive oxygen species (ROS) production evaluation by DAF-FM/DA and DHE fluorescence

In order to analyze the intracellular production of NO and ROS, HAoEC treated with PBS or OSK-SeV (with no EGFP) were incubated with the selective intracellular

NO dye DAF-FM/DA (Invitrogen, D23844; 10 µM) or intracellular ROS dye Dihydroethidium (DHE, Invitrogen, D23107; 10 µM), respectively. The intracellular fluorescence intensity of DAF-FM/DA or nuclear fluorescence intensity (A.U.) of DHE was evaluated.

### LVCO and LV-OSK treatment in mice

One intravenous (i.v.) injection of LV carrying control plasmid (LVCO) or LV containing the transcription factors Oct-3/4, Sox-2 and Klf-4 (LV-OSK) was performed in male and female BPN/3J and BPH/2J (40-44-week-old) mice through the tail vein. One separate experiment involved one intraperitoneal (i.p.) injection of 100 µL (5 µL LVCO or LV-OSK + 95 µL sterile saline) only in female BPN/3J and BPH/2J mice at 30 weeks of age. Regardless of the route of administration, mice were placed in their respective cages and monitored until recovered, and experiments were performed 10 days post-treatment.

### Echocardiography in mice

Male and female BPH/2J and BPN/3J mice, i.v. treated with LVCO or LV-OSK and at 40-44 weeks of age had their cardiac function measured by echocardiography.

### Blood pressure measurement by left carotid catheterization

Treated male and female BPN/3J or BPH/2J mice were anesthetized with 3% isoflurane (1 L/min 100% oxygen) and placed in the supine position on a warm pad. A small incision on the skin was made on the left side of the neck, and the left carotid artery was catheterized. The isoflurane anesthesia was reduced by 1%, and the pulsatile arterial pressure (PAP, in mmHg) was acquired for 20 minutes after the stabilization of the signal, using the LabChart 7 Software. After blood pressure measurements, blood was collected through the arterial catheter in chilled heparinized tubes under anesthesia (5% isoflurane). Plasma was obtained after centrifugation at 1,000 x g for 15 min at 4°C, and stored at -80°C until experiments. The systolic blood pressure (SBP, in mmHg) was calculated and analyzed using the LabChart 7 Software formulas.

### Vascular function in mesenteric resistance arteries (MRA)

Second-order MRA (with an inner diameter up to 250 µm) were isolated from male and female LVCO- or LV-OSK-treated BPN/3J and BPH/2J mice (i.v.), and 2 mm length segments were mounted on DMT wire myographs (Danish MyoTech, Aarhus, Denmark). The MRA was oxygenated (95% O_2_ and 5% CO_2_), heated (37°C) and submerged in regular Krebs solution. The MRA were normalized to their optimal lumen diameter for active tension development. To test vascular smooth muscle cell integrity, the arteries were initially contracted with modified 120 mM high potassium chloride (KCl) Krebs solution. Cumulative concentration-effect curves to ACh (1 pM to 10 μM) after a previous contraction elicited by the thromboxane A2 receptor analogue U46619 (30 nmol/L) was performed to evaluate vascular relaxation. Cumulative concentration-effect curve to the selective α1-adrenergic receptor agonist phenylephrine (PE; 0.1 nmol/L to 100 µmol/L) or to U46619 (1 pmol/L to 1 µmol/L) were also performed, and values were represented as milliNewton per millimeter (mN/mm).

### Plasma TNF-α

Plasma TNF-α was measured in plasma samples obtained from male and female BPN/3J and BPH/2J LVCO- or LV-OSK-treated mice (i.v.), and processed with Mouse TNF-α high sensitivity ELISA kit (eBioscience, BMS607HS), according to the manufacturer’s instructions.

### Plasma estradiol

The estrogen levels in plasma samples were determined in female mice (40-44-week-old) i.v. treated with LVCO or LV-OSK using a commercial ELISA kit (Cayman Chemical, #501890), according to the manufacturer’s instructions.

### Transmission electron microscopy (TEM)

Cell and vascular morphologies were evaluated in mouse ECs or MRA. Briefly, mouse ECs treated with PBS, LVCO or LV-OSK for 3 days, or freshly dissected male mouse MRA i.v. treated with LVCO or LV-OSK (40-44 weeks of age) were processed according to the transmission electron microscopy (TEM) protocol, and images were acquired with a JEOL 1400 Plus Transmission Electron Microscope.

### Multi-photon microscopy and second harmonic generation (SHG)

MRA isolated from male LVCO- or LV-OSK-i.v. treated BPN/3J and BPH/2J mice (at 40-44 weeks of age) was investigated on their collagen, elastin content, and EGFP signals by second harmonic generation (SHG) signals using multi-photon microscopy.

### Statistical analysis

All statistical analyses were performed using GraphPad Prism 9.0 (GraphPad Software Inc., La Jolla, CA, USA). Data are presented as mean ± standard error of the mean (S.E.M) and statistical significance was set at *p*<0.05. All bar graphs are accompanied by dispersion of the individual values (n). The unpaired and two-tailed Student’s t test was used to compare two different groups, or the One-way or two-way analysis of variance (ANOVA), followed by Tukey post hoc was used to identify interaction factors and compare 3 or more groups.

## Results

### Mouse EC *In Vitro* Reprogramming by Overexpression of Oct-3/4, Sox-2, and Klf-4 (OSK) Transcription Factors

The generation of pluripotent stem cells from cultured fibroblasts by the incorporation of four transcription factors Oct-4, Sox-2, Klf-4, and c-Myc, collectively called the “Yamanaka” or “OSKM factors” [11], changed the progression of cellular rejuvenation research. However, c-Myc is an oncogene associated with chromosomal instability and risk of tumorigenesis constitutively expressed in over 70% of human cancers [14]. Recently, it was shown that the ectopic activation of three Yamanaka factors, Oct-4, Sox-2, and Klf-4 (OSK) delivered by the widely used adeno-associated viral (AAV) vectors in mouse retinal ganglion cells in a mouse model of glaucoma and aged mice restored the youthful DNA methylation patterns and transcriptomes, promoting axon regeneration and vision loss reversion [12].

We selected a lentiviral vector (LV) to deliver the OSK plasmid due to its high capacity for assembling >10 kb sequences and to ability to integrate into nondividing cells [15]. We used a LV carrying the enhanced green fluorescence protein (EGFP) as a reporter and the EC-specific cadherin 5 (Cdh5) promoter with more than 12 kb sequences (Fig. 1 A). The three OSK transcription factors were forcefully expressed in mouse primary vascular ECs (Fig. 1 and 2). The control group is LV-EGFP-Cdh5 without OSK factors (referred to as LVCO from here on), while the treatment group is the LV-EGFP-Cdh5 with OSK factors (referred to as LV-OSK from here on). We observed EGFP^+^ ECs in both LVCO and LV-OSK groups (Fig. 1 B, F, and G). The estimated transfection efficiency was 44.55 ± 1.66% for the LVCO, and 19.04 ± 0.85% (n=5) for the LV-OSK. No differences in EC morphology and intracellular organelles (Fig. 1C), or induction of apoptosis and necrosis were evident after treatments (Fig. 1 E). The integrity of the LV-OSK was confirmed by transmission electron microscopy (TEM) (Fig. 1D).

**Fig. 1.**
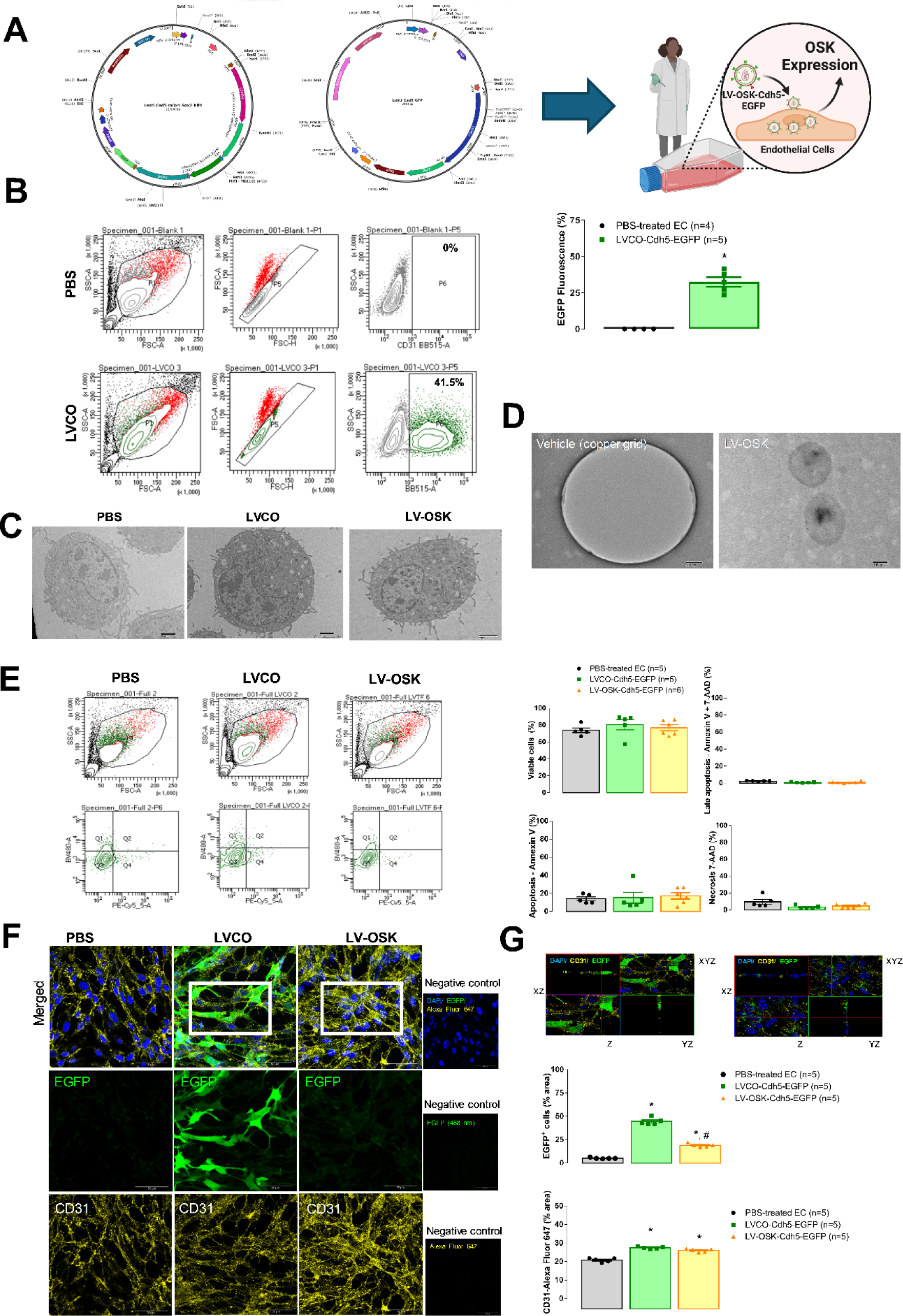
Mouse EC *in vitro* reprogramming with increased levels of EGFP^+^ cells. In **A**, LV-OSK (left) and LVCO (right) maps and graphical abstract of experimental design. In **B**, dot plots, and bar graphs show mouse ECs presented with increased EGFP^+^ cells, after LVCO treatment compared to PBS-treated cells. In **C**, TEM images obtained from PBS-, LVCO- or LV-OSK-treated mouse ECs. In **D**, negative staining for TEM shows the LV-OSK particles. In **E**, LV-OSK, LV-CO or PBS treatment did not induce apoptosis or necrosis in mouse ECs. Dot plots are presented on the left and fluorescence intensity bar graphs are shown on the right. Representative images (**F** and **G**) of mouse ECs treated with PBS, LVCO or LV-OSK. The immunofluorescence assay shows positive staining for DAPI (in blue; nuclei), EGFP (in green) and CD31 (in yellow). Bar represents 50 µm. Negative controls were cells with EGFP detection and secondary antibody incubation with DAPI. The EGFP^+^ cells and CD31 quantification are presented in **G**. Zoomed areas (**G**) from representative images on **C**, showing orthogonal sections (*xz*, *yz*, and *xyz*) and bar represents 10 µm. * Different from PBS-treated cells; # different from LVCO.

**Fig. 2.**
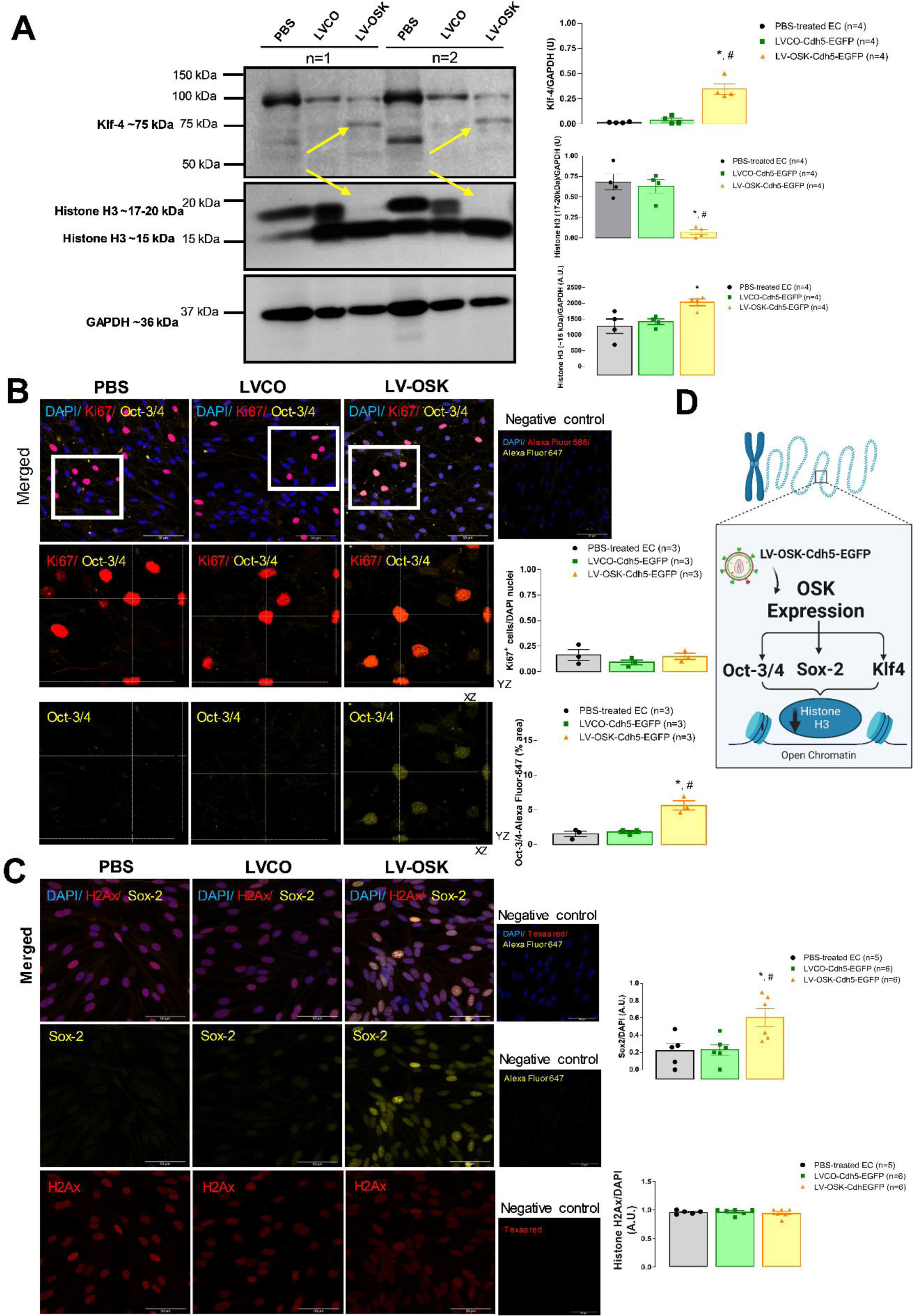
ECs *in vitro* reprogramming induced Oct-3/4, Sox-2 and Klf-4 overexpression and modulation of histone H3 levels. In **A**, representative Western blotting membrane (left) for Klf-4 (monomeric form at ∼75 kDa yellow arrows), and total histone H3 (upper band at 17-20 kDa, and lower band at ∼15 kDa. The Klf-4 and histone H3 levels are presented in bar graphs (right). In **B**, representative images of mouse ECs show the overexpression of Oct-3/4 (yellow) in LV-OSK-treated cells. Upper panel with merged images shows DAPI (blue), Ki67 (red) and Oct-3/4 (yellow). Bar represents 50 µm. Middle and lower panels represent the zoomed (161.05%) orthogonal sections (*xz*, *yz* and *xyz*) from the upper panel for Ki67^+^ and Oct-3/4^+^ cells (middle) or Oct-3/4^+^ cells (lower). Bar graphs present the quantification of Oct-3/4 and Ki67/DAPI ratio from the upper panel images. In **C**, representative images of mouse ECs show the overexpression of Sox-2 (yellow) in LV-OSK-treated cells. Upper panel with merged images showing DAPI (blue), total histone H2Ax (red) and Sox-2 (yellow). Bar represents 50 µm. Negative controls were cells with secondary antibody and DAPI. Middle and lower panels show Sox-2^+^ cells (yellow) or total histone H2Ax (red) in separated images from upper panel, respectively. Bar graphs show the quantification of Sox-2/DAPI nuclei and total histone H2Ax/DAPI ratio from the upper panel images. In **D**, graphical abstract illustrates LV-OSK effects on chromatin remodeling in ECs. * Different from PBS-treated cells; # different from LVCO-treated cells.

We observed that the EC marker cluster of differentiation 31 (CD31), also referred to as platelet endothelial cell adhesion molecule 1 (PECAM1), increased in both LV-treated ECs compared to the cells treated with phosphate-buffered saline (PBS) (Fig. 1 G). The monomeric form of Klf-4 (Fig. 2 A; yellow arrow; ∼75 kDa, as described by Mastej et al. [16]) was upregulated only in LV-OSK-treated cells. Along with Klf-4 modulation, we observed the presence of histone H3 proteins (Fig. 2 A; native H3, 15 kDa and 17-20 kDa). Interestingly, the upper band (17-20 kDa) was almost undetectable in the LV-OSK-treated ECs only, suggesting that OSK-induced modification of histone H3 could be a crucial factor in the EC reprogramming *in vitro* (Fig. 2 A and D). Next, we observed Oct-3/4 and Sox-2 upregulation in the LV-OSK-treated ECs (Fig. 2 B and C, respectively), but not in controls. There were no differences in cellular proliferation which was evidenced by Ki67 staining (Fig. 2 B). In regard to nuclei function, histone H2Ax is a variant found in almost all eukaryotes. It contributes to genome stability by laying the foundation for the assembly of repair foci to counteract DNA damage [17]. Here, we observed similar H2Ax levels in PBS or both LV-treated ECs, suggesting an intact nuclei integrity and function upon LV infections (Fig. 2 C).

### Overexpression of OSK Induced Endothelial Progenitor Cell (EPC)-Like Features and Prevent EndMT *In Vitro*

The CD133 is an EPC marker, which is highly expressed on immature stem cells but is absent in mature ECs after differentiation [18,19]. EPCs are a heterogeneous group of cells characterized by the expression of surface markers CD133^+^/CD34^+^/VEGFR2^+^ for a more primitive population [20]. We observed a significant increase in the CD133^+^ ECs in the LV-OSK group than the other groups (Fig. 3 A). However, there was no significant difference in the CD34^+^ cells among the groups and there was no positive staining for CD45, a general marker for leukocytes (Fig. S1 B). In addition, increased EC migration is a hallmark of EndMT [21]. Interestingly, the ECs treated with LV-OSK decreased migration and senescence-associated (SA) β-galactosidase activity (Fig. 3 B and C), suggesting a protective effect of OSK against cell senescence. Furthermore, we observed that ECs treated with LVCO presented some features of EndMT with an increased level of α-smooth muscle actin (α-SMA) (Fig. 3 D). However, this phenomenon was not observed in the LV-OSK-treated ECs, suggesting an EndMT-counteractive role of OSK (Fig. 3 E).

**Fig. 3.**
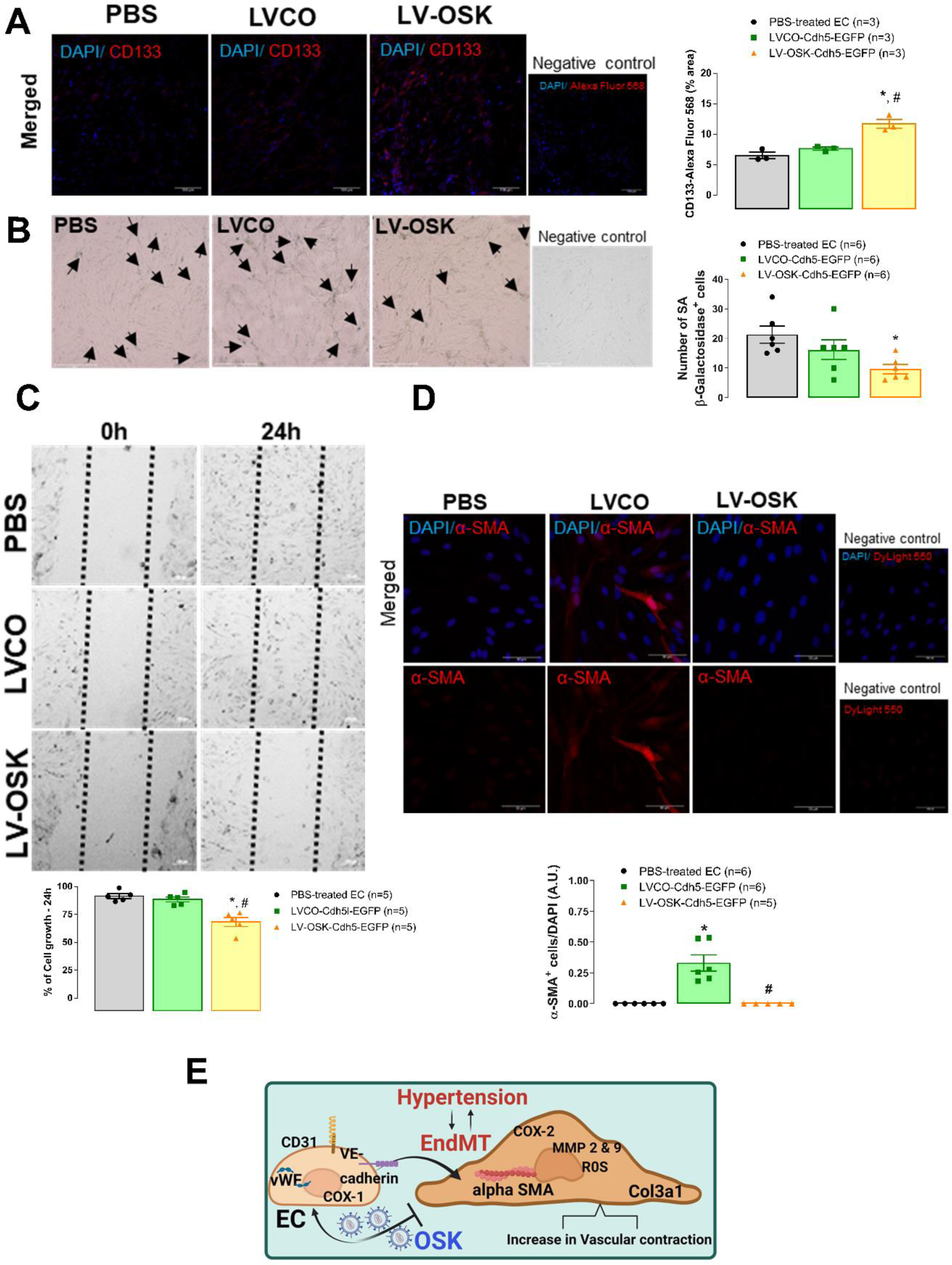
EC *in vitro* reprogramming induced an EPC-like phenotype that prevented cell migration, senescence and EndMT. In **A**, representative images and bar graph show the quantification of mouse ECs CD133 upregulation (red) in LV-OSK-treated cells. Negative control represents cells with secondary antibody and DAPI (blue). Images were acquired on confocal microscope and bar represents 100 µm. In **B**, SA β-galactosidase assay in mouse ECs representative images demonstrate positive blue staining (arrows) for the senescence marker. Bar graph shows the number of positive blue cells for SA β-galactosidase assay. Negative control shows cells with no treatment. Images were acquired on a stereo microscope and bar represents 200 µm. In **C**, wound healing assay shows reduced cellular growth after LV-OSK treatment. Dotted lines denote % cellular growth in 24 h and values are expressed in the bar graph. In **D**, representative images and quantification of mouse ECs show positive staining for α-SMA (red) only in LVCO-treated cells, suggesting EndMT. The upper panel shows merged images for DAPI (blue) and α-SMA (red), and lower panel shows only α-SMA images. Negative control represents ECs with secondary antibody and DAPI. Images were acquired on confocal microscope and bar represents 100 µm. In **E**, graphical abstract illustrates LV-OSK preventing hypertension-induced EndMT in ECs. * Different from PBS-treated cells; # different from LVCO-treated cells.

### *In Vivo* Detection of EGFP and EndMT Was Mitigated by *In Vivo* Reprogramming of ECs in Hypertensive Mice

Mounting evidence suggests that aberrant EndMT is involved in cardiovascular diseases [3,22,23,13]. One of the most potent inducers of EndMT is tumor necrosis factor alpha (TNF-α) [21,24]. We treated mice ECs with TNF-α (Fig. S1 A), and observed a downregulation of EC-associated genes and an upregulation of EndMT-associated gene expression. Specifically, the levels of *Pecam-1*, *vascular endothelial* (*VE)-cadherin*, *eNOS* (*NOS3*), *COX-1*, and *Von Willebrand factor* (*vWF*) genes were significantly reduced and the levels of metalloproteinases (*MMP*)-*2* and *9*, *Acta2* (*α-SMA*), and *collagen* (*Col3a1*) genes were upregulated in ECs treated with TNF-α.

Next, we used an inbred mouse model of spontaneous hypertension, the BPH/2J mouse strain [25] to understand whether EndMT also occurs in essential hypertension. In addition, to understand whether ECs reprogramming would attenuate EndMT formation in hypertensive animals, we infected normotensive (BPN/3J) and hypertensive animals (BPH/2J) with LVCO and LV-OSK for 10 days (single-short partial reprogramming), and we checked for EGFP detection. As shown in Figure 4 A, both treatment with LVCO or LV-OSK showed EGFP signals (in green) in the prefrontal cortex of male and female BPN/3J and BPH/2J mice. Furthermore, it was observed that CD31 expression (in red) was high in normotensive males treated with LV-OSK, but not in hypertensive males or in the female group, suggesting that the single-short partial reprogramming of ECs induced an angiogenic effect in normotensive males, but not in female mice. An impaired response to the OSK-induced angiogenesis was observed in both male and female hypertensive groups. Additionally, we observed an exacerbated formation of EndMT in the coronary artery (Fig. 4 B and C), mesenteric resistance artery (MRA) (Fig. 4 D), and thoracic aorta (Fig. 4 E and F) from hypertensive mice (BPH/2J). To the best of our knowledge, this is the first evidence showing that BPH/2J hypertensive mice exhibit EndMT.

**Fig. 4.**
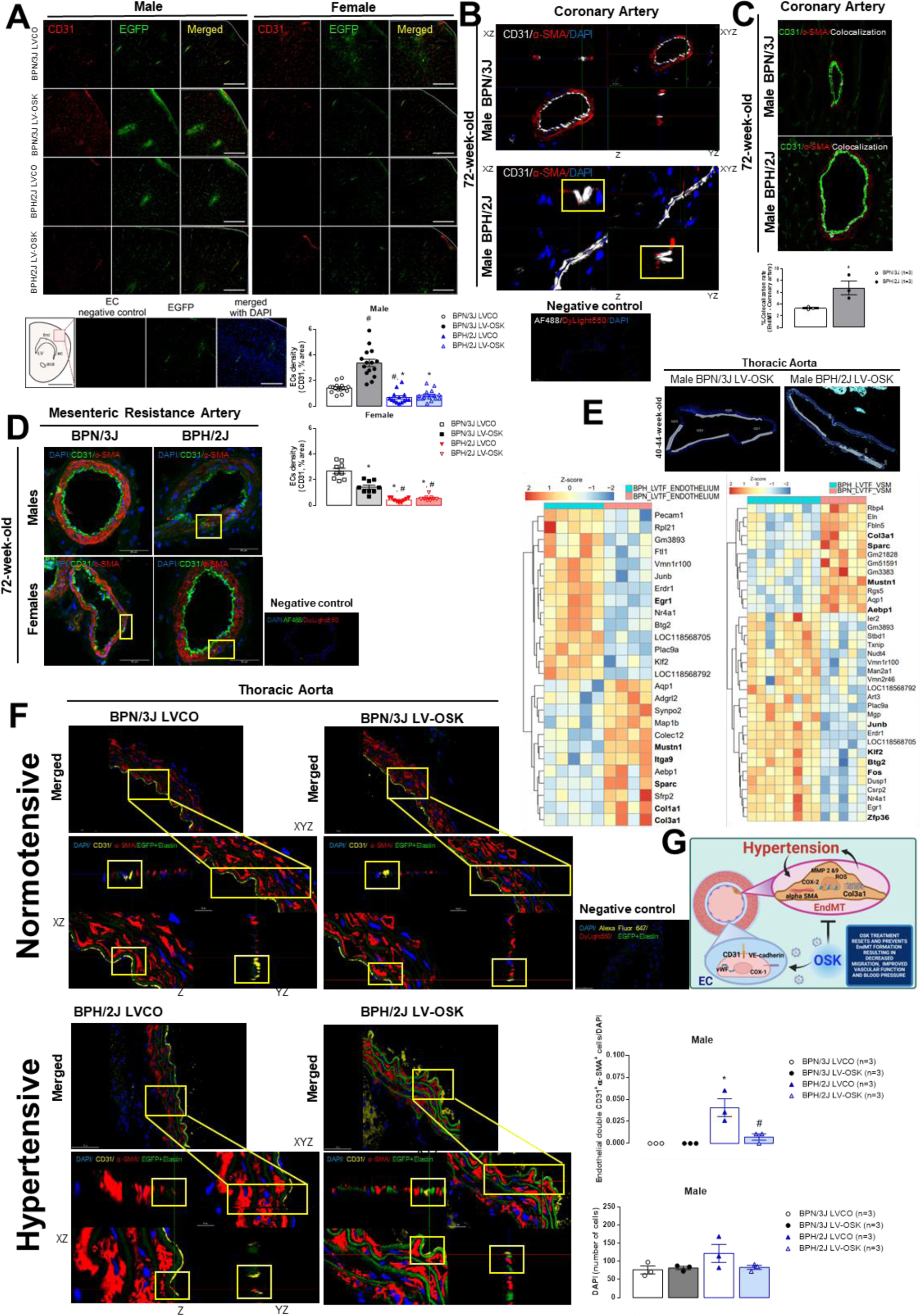
*In vivo* EGFP detection, and EndMT was present in arteries from male and female BPH/2J mice and OSK *in vivo* treatment prevented EC phenotypic transition. In **A**, confocal maximum projection images show immunofluorescence for CD31 (red) and EGFP (green) from the respective LV treatments, in the prefrontal cortex of BPN/3J and BPH/2J male and female mice treated with LV-OSK or LVCO. The summary data indicates the endothelial cell density (CD31) percentage occupied within the area (dotted lines within merged images) in the prefrontal cortex across all groups. The diagram (lower panel) shows the region where all images were taken within the prefrontal cortex. Representative photomicrographs of the negative control for endothelial cells (CD31 immunofluorescence), taken in series with the EGFP (readily fluorescent at 488nm excitation) and DAPI, respectively, in a normotensive male EGFP-SeV-OSK mice. No signal was observed in the CD31 channel with the omission of the primary antibody (Scale bar: 200 µm). *Fml*: forceps minor of the corpus callosum; *LV*: lateral ventricle; *ec*: external capsule; *ACA*: anterior commissure, anter part. Scales: 2 inches (diagram) and 200 µm (photomicrograph). The diagram is based on Figure 19, with bregma 1.41 mm from Paxinos and Franklins [65]. Data were analyzed using one-way ANOVA with Tukey’s post-hoc analysis (*p*<0.05). ^#^ different from BPN/3J LVCO; * different from BPN/3J LV-OSK. In **B**, representative images of coronary artery isolated from a 72-week-old male BPH/2J mice (lower panel) with no treatments evidencing EndMT by double positive ECs (yellow rectangles) to CD31 (white) and α-SMA (red) compared to an age-matched male BPN/3J mice (upper panel). Orthogonal images (*xz*, *z*, *yz*, and *xyz*) highlight the double positive CD31^+^ and α-SMA^+^ ECs suggesting the hypertension-induced EndMT. Negative control represents artery with secondary antibodies and DAPI. Images were acquired on confocal microscope and bar represents 10 µm. In **C**, representative images (upper panel) and bar graph (lower panel) with the quantification of the % colocalization (white) of double positive CD31 (green) and α-SMA (red) signals in coronary arteries from male BPN/3J and BPH/2J mice at 72 weeks of age with no treatments. Unpaired Student’s *t* test (*p*<0.05). * different from BPN/3J mice. In **D**, representative images of double positive CD31 (green), α-SMA (red) and DAPI (blue) in MRA isolated from male (upper panels) or female (lower panel) BPN/3J and BPH/2J mice at 72 weeks of age with no treatments. Bar represents 50 µm, and negative control represents artery with secondary antibodies and DAPI. In **E**, GeoMx analysis of thoracic aortas isolated from a 40-44-week-old male BPN/3J or BPH/2J mice treated with i.v. injection of LV-OSK showing region of interest (ROI) selected in the endothelial or vascular smooth muscle (VSM) layers. Heatmaps show EndMT genes in ECs (left panel) or VSM (right panel). In **F**, representative EndMT images of thoracic aortas from male BPN/3J and BPH/2J mice (40-44-week-old) i.v. treated with LVCO (left panel) or LV-OSK (right panel). Upper images and lower images are the orthogonal zoomed (2x) sections (*xz*, *yz* and *xyz*; bar represents 20 µm) to show double positive CD31^+^ and α-SMA^+^ cells in the vascular endothelium of BPH/2J-LVCO mice, but not in BPH/2J LV-OSK-treated or BPN/3J mice (yellow rectangles). Merged images show DAPI (blue), CD31 (yellow), α-SMA (red) and elastin+EGFP^+^ cells (green), and bar represents 50 µm (upper panel). EGFP^+^ signals were derived from LVCO or LV-OSK treatments. Negative control represents aortas with secondary antibodies, elastin autofluorescence and EGFP detection with DAPI. Bar graphs show the number of double positive CD31^+^ and α-SMA^+^ cells to DAPI ratio in the aortic endothelium, and the total number of vascular DAPI^+^ nuclei, respectively. Data were analyzed using two-way ANOVA with Tukey’s post-hoc analysis (p<0.05). * Different from BPN/3J LVCO; # different from BPH/2J LVCO. In **G**, graphical abstract summarizing hypertension induced EndMT, and the OSK strategy to promote ECs *in vivo* reprogramming to reverse EndMT.

GeoMx digital spatial profile, is a robust spatial multi-omic platform for analysis of profile expression of RNA and protein from distinct tissue compartments. Through this platform, we observed that LV-OSK treatment increased levels of EC function-related genes such as *Pecam-1* and *Klf-2*, and reduced levels of EndMT-related genes such as *Col1a1* and *Col3a1* in the endothelium from BPH/2J mice (Fig. 4 E). LV-OSK treatment also decreased *Col3a1* gene in the vascular smooth muscle (VSM) layer of BPH/2J mice (Fig. 4 E). Additionally, the amount of double CD31^+^ and α-SMA^+^ cells was reduced in the endothelium of LV-OSK-treated BPH/2J mice (Fig. 4 F), with no differences in the total number of cell nuclei (Fig. 4 F) or other parameters (Fig. S5 A-C), suggesting the prevention of EndMT induced by LV-OSK. A graphical abstract illustrates the OSK prevention of hypertension-induced EndMT (Fig. 4 G).

### Reprogramming ECs *In Vivo* Decreased Blood Pressure and Restored Vascular Function in Hypertensive Mice

Here, for the first time, we observed that *in vivo* treatment with LV-OSK decreased blood pressure in male BPH/2J mice at 40-44 weeks of age (Fig 5A and B). There were no effects induced by LVCO or LV-OSK treatments on heart rate, and body weight/tibia length ratio in male or female BPN/3J mice (Fig. 5 B-D). No death occurred during the treatments. The LVCO or LV-OSK treatment did not induce lung edema or improve plasma TNF-α levels (Fig. S3). Surprisingly, LV-OSK did not modify female BPH/2J mice blood pressure levels at 40-44 weeks of age (Fig. 5 B), and for this reason, we decided to treat female BPH/2J mice at an early age (30-week-old). We observed that OSK treatment in female BPH/2J mice at an early age improved the blood pressure values, since no more differences were observed between female normotensive LV-OSK *vs*. hypertensive LV-OSK mice at 30-week-old (Fig. S2), suggesting that female hypertensive mice are more susceptible to EC reprogramming at an early age. No change in estrogen levels was observed in female mice (Fig. 5 E). As expected, cardiac dysfunction was present in the hypertensive animals at 40-44 weeks of age. However, the treatment did not improve these parameters (Fig. S4 and Supplemental Table S1). On the other hand, LV-OSK treatment improved vascular hypercontractility in resistance arteries from male hypertensive mice (40-44-week-old) regardless of the agonist used (Fig. 6 A-D). The treatment did not change vascular function in female mice (40-44 weeks of age). Further, no changes were observed in the lumen diameter from the MRA in male mice treated with LV-OSK (Fig. S3 C). Since we observed an improvement in EndMT and vascular hypercontractility in arteries from male mice treated with LV-OSK, we then investigated whether it could be due to an attenuation of elastin breaks, since EndMT increases MMPs (Fig. S1 A). Accordingly, LV-OSK alleviated resistance arteries elastin breaks in the internal elastic lamina (IEL) in BPH/2J mice compared to hypertensive LVCO-treated mice (Fig. 6 E). No elastin breaks were detected in arteries from BPN/3J mice (Fig. 6 E). No differences were observed in MRA collagen total area, but LV-OSK increased EGFP^+^ and elastin signals in MRA from male BPH/2J mice (Fig. S5 D).

**Fig. 5.**
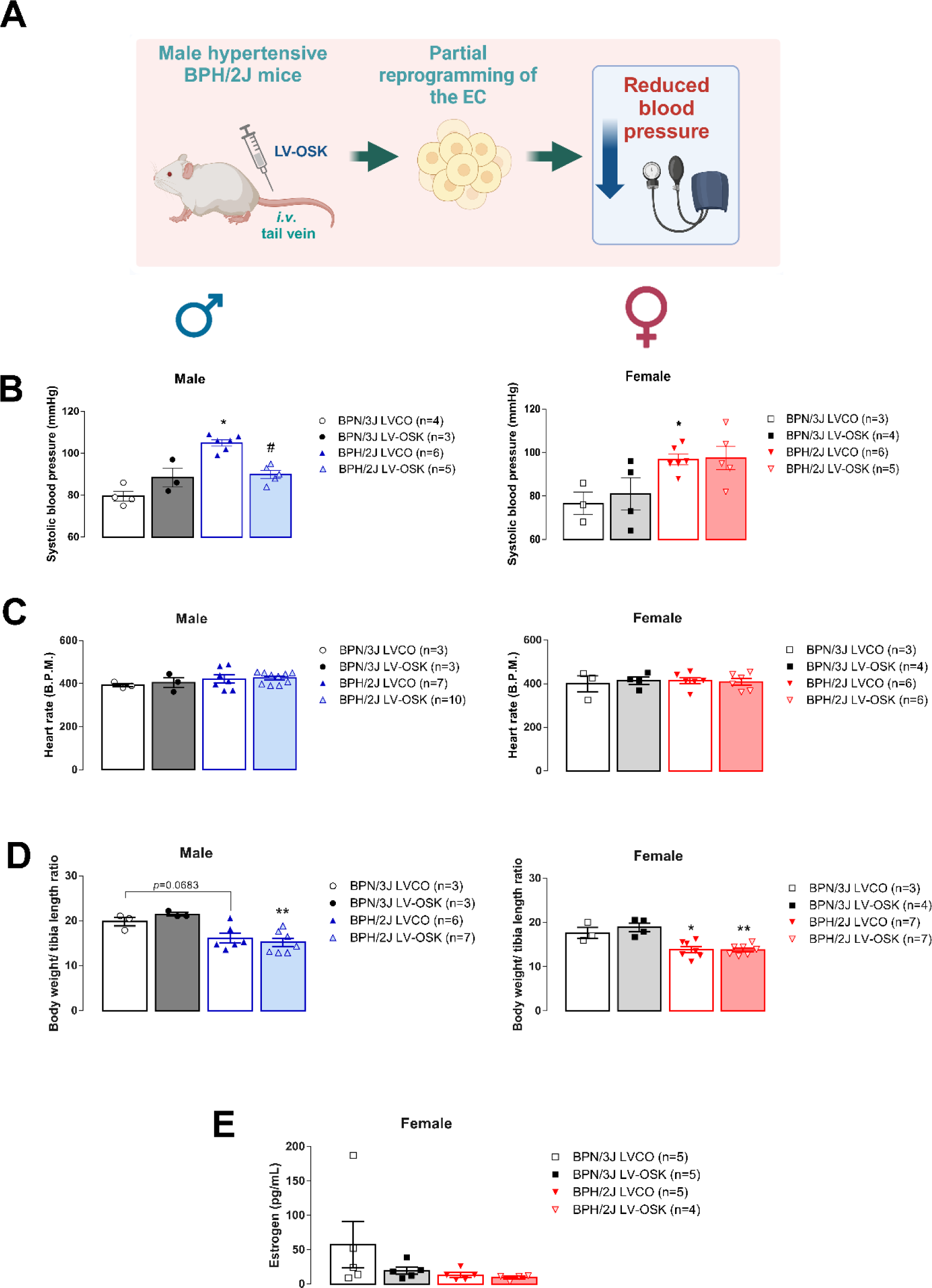
ECs *in vivo* reprogramming reduced blood pressure in male hypertensive BPH/2J mice. In **A**, graphical abstract shows the LV-OSK *in vivo* treatment in mice. **B** and **C** represent the systolic blood pressure values and heart rate, respectively measured from male and female BPN/3J and BPH/2J mice (40-44-week-old), treated with LVCO or LV-OSK (i.v.) at 10 days post-treatment. In **D**, bar graphs represent the body weight/tibia length ratio from male and female mice, respectively. In **E**, estrogen levels measured in plasma samples from female BPN/3J and BPH/2J mice (40-44-week-old) i.v. treated with LVCO or LV-OSK. * Different from BPN/3J LVCO; # different from BPH/2J LVCO; ** different from BPN/3J LV-OSK.

**Fig. 6.**
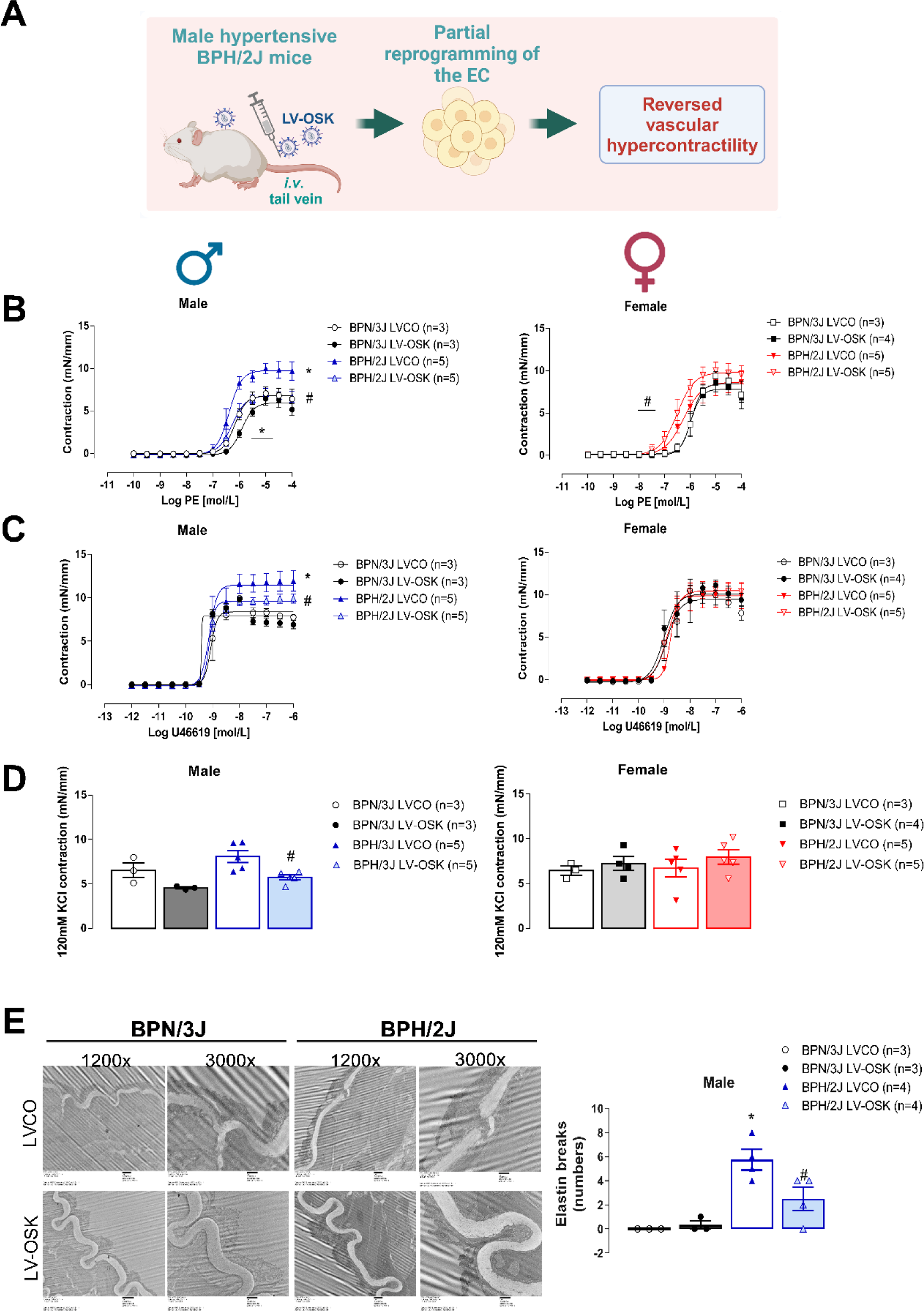
MRA hypercontractility is reversed by EC *in vivo* reprogramming in male BPH/2J mice. In **A**, graphical abstract shows LV-OSK treatment reversing vascular hypercontractility in hypertensive male mice. Vascular reactivity assay showing cumulative concentration-effect curves to phenylephrine (PE) or to U464619 in MRA isolated from male (left) or female (right) BPN/3J and BPH/2J mice, treated with LVCO or LV-OSK (i.v., at 40-44 weeks of age) (**B** and **C**). *In bolus* MRA contraction induced by high potassium Krebs solution (**D**). In **E**, TEM images obtained after vascular reactivity assays on MRA isolated from male BPN/3 and BPH/2J mice (40-44 weeks of age), treated with LVCO or LV-OSK (i.v.). MRA images highlight internal elastin lamina (IEL) and the intact vascular endothelium. Bar graph shows the number of elastin breaks in MRA IEL. * Different from BPN/3J LVCO; # different from BPH/2J LVCO; ** different from BPN/3J LV-OSK.

### OSK Treatment Improved Endothelial Function by Increasing the Synthesis of NO and Decreasing ROS Generation in Human Male and Female ECs From Hypertensive Patients

Since EC reprogramming presents as a novel therapeutic approach that might reduce the morbidity and mortality associated with hypertension, we opted to utilize primary HAoECs obtained from normotensive and hypertensive women and men patients. Additionally, we used the Sendai virus (SeV), an RNA virus with no DNA intermediate and no nuclear phase in its lifecycle (Fig. 7 A), to reprogram human ECs [26]. This eliminates the risk of unwanted integration, making SeV a safe option. For mechanistic insights related to EC function, OSK overexpression with SeV (OSK-SeV) in HAoECs restored acetylcholine (ACh, 1 μM)-induced NO biosynthesis; this was further supported by an upregulation of p-eNOS (Ser1177) in OSK-overexpressed HAoECs from hypertensive patients (Fig. 7 B, D, and E). Total eNOS expression did not change (Fig. S1 C), and CD133 expression representative images showed a trend to increase after OSK-SeV (Fig. S1 D). It is important to note that OSK-SeV did not induce α-SMA expression (Fig. S1 D). In addition, we observed that OSK expression reduced intracellular ROS generation but not in EC from normotensive patients (Fig. 7 C). These data demonstrate that OSK overexpression can improve the physiological function of HAoECs from hypertensive patients.

**Fig. 7.**
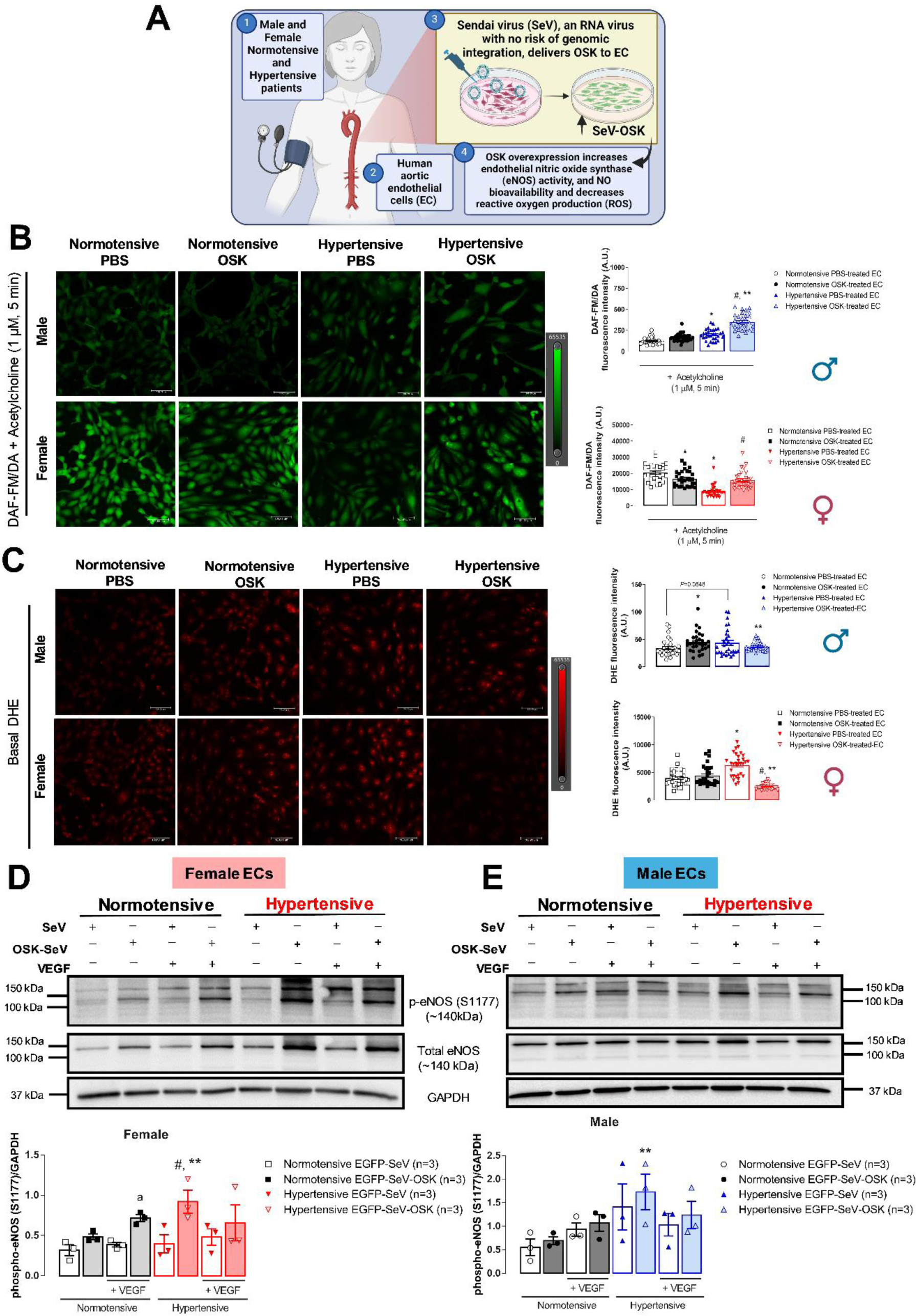
OSK overexpression in human ECs from hypertensive patients restored eNOS activation, NO production and reduced reactive oxygen species (ROS) production. In **A**, graphical abstract illustrating the mechanism of action of OSK in human aortic ECs from hypertensive patients (HAoECs). In **B**, intracellular NO production in male (upper panel and bar graph) and female (lower panel and bar graph) ECs isolated from normotensive and hypertensive patients, treated with PBS or with the overexpression of OSK using SeV. HAoECs were stimulated with acetylcholine, and NO production was measured by DAF-FM/DA (green). Images were acquired on confocal microscope and bar represents 100 µm. In **C**, intracellular ROS production in male (upper panel and bar graph) and female (lower panel and bar graph) ECs from normotensive and hypertensive subjects, treated with PBS or with SeV-OSK in basal condition. ROS production was measured by DHE. Images were acquired on confocal microscope and bar represents 100 µm. * Different from normotensive PBS-treated cells; # different from hypertensive PBS-treated cells; ** different from normotensive OSK-treated cells. **D** and **E** show Western blotting representative membranes and bar graphs of phospho-eNOS (Ser1177; upper), total eNOS (middle) and GAPDH (lower), in basal (-) or after (+) VEGF stimulation from female (**D**) and male (**E**) ECs from normotensive and hypertensive patients, treated with SeV-EGFP or EGFP-OSK-SeV. GAPDH was used as loading control. # Different from hypertensive EGFP-SeV-treated cells in basal condition; ** different from normotensive EGFP-OSK-SeV-treated cells in basal condition; ^a^ different from normotensive EGFP-SeV-treated cells+VEGF.

We next wanted to see whether the counteractive function of OSK against EndMT in hypertensive mice could be recapitulated in human ECs from hypertensive patients. HAoECs were infected by SeV expressing OSK or EGFP reporter genes. The cells were collected at day 0 (mock), 3, or 7 post-infection, followed by a single-cell RNA sequencing analysis. Upon dimensional reduction and Leiden clustering, the cells were assigned into 12 clusters based on the single-cell transcriptomes (Fig. 8 A). The time points did not show any enrichment among clusters (Fig. 8 B). Samples from different sexes were also evenly distributed among clusters (Fig. 8 C). In addition, the expression of OSK factors was mainly in cluster 12 (Fig. 8 D). ECs in clusters 3 and 9 showed an elevated basal level of KLF4 with low levels of SOX2 and POU5F1 (the human Oct-4 gene) than ECs in other clusters, except cluster 12 (Fig. 8 D). We next looked into the representative differential gene expression (DEGs) among clusters 3, 9, and 12 (Fig. 8 E). Cluster 3 showed high levels of EC markers such as TEK, CDH5, KDR, and PECAM1 but also exhibited elevated levels of EC senescent markers such as ESM1 [27,28], MMP1 and *2* [29], CDKN1a and *2a* [30,28], IFGBP5 [31,32], IFI27, IFIT1, and PLAT [32]. These data suggest that cluster 3 mainly represents the senescent ECs (Fig. 8 E). The ECs in cluster 9 showed the absence of typical EC markers but instead had high levels of EndMT-related markers such as CNN2, COL1a1, LAMA2, IGFBP3, PDGFRA, ITGA4, COL3a1, suggesting a subgroup of EndMT-induced ECs (Fig. 8 E). The ECs in cluster 12, representing the OSK subgroup, showed lower levels of EC senescent markers than the ECs in cluster 3, lower levels of EndMT markers than the ECs in cluster 9, and higher levels of EC functional-related genes such as NOS3, KLF3, IGFBP6, IFG2, and KLF4 than the cells in clusters 3 and 9 (Fig. 8 E). The EC markers (TEK, CDH5, KDR, and PECAM1) in the cells of cluster 12 showed intermediate levels between the cells in clusters 3 and 9 (Fig. 8 E), indicating a process of regaining EC phenotypes (partial EC reprogramming). This pattern was further shown in the EC from the OSK subset *versus* the GFP subset, from cluster 12 *versus* cluster 3 (Fig. 8 F) or cluster 9 (Fig. 8 G) in both male and female ECs from hypertensive patients. These data demonstrated that OSK alleviated EC senescence and mitigated EndMT, facilitating the restoration of EC phenotypes in human ECs from hypertensive patients.

**Fig. 8.**
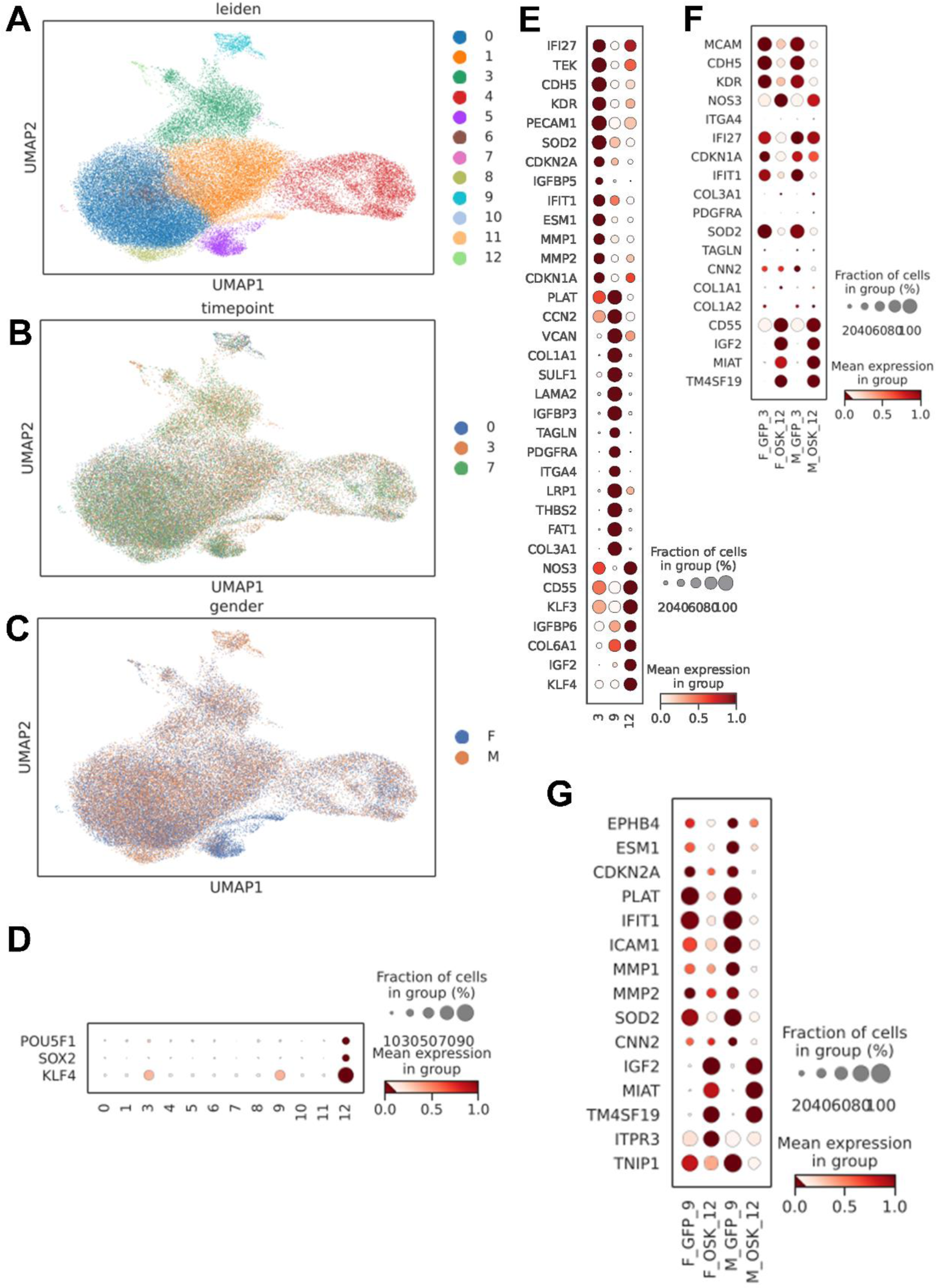
SeV-delivered OSK alleviated EC senescence and mitigated EndMT in human ECs from hypertensive patients. In **A**, UMAP plot of the single-cell transcriptomes for 12 cell clusters. **B** and **C**, treatment time points and sample genders were not contributors to cluster formation. **D**, dot plot shows that OSK genes are mainly expressed in the cells of cluster 12. **E**, dot plot shows the representative DEGs of each cell cluster of 3, 9, and 12. **F** and **G**, dot plot shows the representative biomarkers in subsets of GFP and OSK from cluster 12 *versus* cluster 3 (**F**) or cluster 9 (**G**) among both male and female ECs from hypertensive patients. M, male; F, female; GFP, green fluorescent protein (SeV control expressing GFP only).

## Discussion

According to the 2017 Guideline for the Prevention, Detection, Evaluation, and Management of High Blood Pressure in Adults [33], 130/80 mmHg rather than 140/90 mmHg is considered stage 1 hypertension. Nonetheless, the American Heart Association (AHA) in conjunction with the National Institutes of Health (NIH), annually reports the most up-to-date statistics related to hypertension, and based on the 2024 Statistical Update [34], the age-adjusted prevalence of hypertension among US adults at ≥20 years of age was estimated to be 46.7% (50.4% for men and 43.0% for women). However, for those at ≥65 years of age, the percentage of women with hypertension was greater [34]. Therefore, there is a consensus from the scientific community that efforts should not only consider specific research for the early detection, monitoring, and treatment of hypertension but also focus on basic discovery and translational research to identify the cause and prevent hypertension and its associated side effects. Here, we demonstrated, for the first time, that ECs reprogramming represents a highly potential and innovative strategy to prevent and reverse hypertension-induced vascular damage. Our scientific approach was based on the overexpression of the three OSK transcription factors in an attempt to bring ECs back to a youthful phenotype in hypertensive mice. Correspondingly, we generated ECs with EPC-like features *in vitro* that expressed lower migratory capacity and prevented cellular senescence. Human ECs from hypertensive patients treated with OSK recovered their ability to activate eNOS and biosynthesize NO, with low ROS levels. In addition, OSK treatment of BPH/2J mice *in vivo* was advantageous, since blood pressure values and resistance arteries hypercontractility were normalized, which contributed to the prevention of EndMT and elastin breaks. The treatment was validated by the *in vivo* EGFP signals in the prefrontal cortex of male and female mice. In addition, LV-OSK did not decrease blood pressure or vascular contraction in resistance arteries from female 40-44-week-old BPH/2J mice. It is known that ∼44-week-old female mice have the endocrine equivalent of human menopause [35]. In this scenario, the cardiovascular system of these mice presents with a fast decline and is less responsive to any treatment [67]. We therefore decided to treat female mice at an earlier age (30 weeks of age) on the grounds that although these female adult mice are hypertensive, they are still capable of reproducing. We observed that early treatment with LV-OSK decreased blood pressure in female mice. These results corroborate the Women’s Health Initiative (WHI) trial studies which showed that early treatment (use of hormone therapy) in younger women (50-59-year-old) or in early postmenopausal women had a beneficial effect on the cardiovascular system. On the other hand, late treatment (>10 years after the establishment of menopause) hormone therapy had some detrimental or minimal beneficial effects [36, 67].

The vasculoprotective action elicited by the pluripotency transcription factors is well documented in the literature. For instance, the Klf-4 is well-known for its anti-inflammatory, anti-adhesive, and antithrombotic activities [37]. It plays a critical role in maintaining ECs quiescence and function [16]. Specific endothelial Klf4-knockout mice exhibit features of EndMT regardless of sex. The pluripotency factor Oct-4 was believed to be dispensable in adult somatic cells. However, Shin and collaborators [38] have demonstrated that Oct-4 plays a functional role in regulating ECs metabolism and phenotypic transition in atherosclerosis. Further, Sox-2 is found to contribute to the reprogramming of human corneal endothelial cells [39], and it plays a role in ECs differentiation from embryonic stem cells, since the endothelial markers emerge between days 3 and 6 of endothelial induction, as the expression of Sox-2 increases [40].

Beyond its established contribution as a transcriptional activator of vasculoprotective genes, Klf-4 is able to act as a chromatin organizer by recruiting the SWI/SNF chromatin remodeling complex to modify chromatin accessibility and control the endothelial enhancer landscape [41]. The mammalian SWI/SNF complex regulates chromatin accessibility, leading to the disruption of histone-DNA contact. Camerini-Otero and Felsenfeld [42] reported the disulfide bond formation on histone H3, and suggested that histone H3 intermolecular disulfides might play a role in further stabilizing nucleosomes that need not be unfolded, perhaps related to transcriptionally inactive regions. Our study showed that OSK-induced reprogramming of ECs was effective in monomerizing histone H3, likely through transcription activation on chromatin.

It has been demonstrated that Oct-4 binds to condensed heterochromatin and promotes its opening [43,44]. Specifically, Oct-4 gradually binds to pluripotency network-related gene loci to activate their expression, often accompanied by H3K27me3, H3K4me3 and H3K27ac modifications [45]. Similarly, Sox-2 binds heterochromatin and facilitates chromatin opening *at loci* containing pluripotency genes at an early reprogramming stage [44]. The combination of Sox-2 and Klf-4 further contributes to the opening of most chromatin during the early stages of reprogramming and promotes the mesenchymal-to-epithelial transition process [44,46]. Moreover, the formation of Oct-4/Sox-2 heterodimers is essential for pluripotency establishment [47].

Depletion of canonical histones results in a more open chromatin structure, defined by a reduced histone-to-DNA ratio. Histone depletion can activate a regulatory mechanism to control DNA repair, transcription, and senescence. The C-terminus region of histone H2Ax can be phosphorylated to generate gamma-H2Ax (γH2Ax), and its dephosphorylation has a half-life of approximately 2h, which is similar to the kinetics of DNA double-strand break repair [48]. Post-translational modifications of histone H2Ax are crucial for function and are related to genomic stability [48,17]. Our study therefore shows that although histone H3 was modulated by OSK treatment, no changes in H2Ax levels were found in our ECs regardless of the treatment, suggesting the lack of degradation of this protein, and that OSK overexpression was unlikely to disturb the genomic stability.

Senescence induced by histone depletion is triggered in response to telomere shortening, which causes the loss of the telomere binding sites and the subsequent relocation of the repressor activator protein 1 (Rap1) from telomeres to the promoters of several senescence genes, contributing to their activation [43]. Consistently, overexpression of core histones prevents senescence, and histone mRNA modulation is reported to increase gradually through early zygotic cell divisions [49], and in differentiating embryonic stem cells [50]. Our OSK treatment successfully prevented senescence in ECs despite modulating the histone H3 levels, suggesting that *in vitro* reprogramming represented a protective mechanism for ECs and is unlikely to promote DNA destabilization. Instead, it prevented the transcription of genes associated with senescence.

Vasoregenerative capacities are reported for EPC. In this context, Friedrich and coauthors [20] showed that CD133^+^ EPC isolated from peripheral blood of healthy humans showed a larger fraction of CD34^+^/CD133^+^/VEGFR2^+^ compared to CD34^-^/CD133^+^/VEGFR2^+^ subpopulation. Upon culturing these cells under conditions enabling endothelial-specific differentiation, CD34^-^/CD133^+^/VEGFR2^+^ cells progressively downregulated surface expression of CD133, while upregulating CD34 and the endothelial marker CD31 [20]. Expression of the stem cell marker CD34 is also found on mature VE-cadherin^+^ ECs [18]. In addition, human coronary atherectomy specimen from stable lesions shows more CD34^+^/CD133^+^ EPC, and mice subjected to carotid artery injury and transplanted with CD34^+^/CD1333^+^/VEGFR2^+^ cells present an attenuation of neointima formation [20]. Additionally, capillary networks in ischemic hindlimbs were observed after human cord blood-derived CD133^+^ progenitors were transplanted into nude mice, followed by augmented neovascularization, and improved ischemic limb salvage [51]. Mouse ECs treated with LV-OSK still showed CD31^+^ and CD34^+^ phenotypes with CD133 upregulation, which could be attributed to partial reprogramming with some vasoregenerative features. In this context, LV-OSK induced an increase in CD31 expression in the prefrontal cortex of male normotensive mice, suggesting an upregulation of angiogenic mechanisms.

It has been demonstrated that in human pulmonary hypertension, the pathological vascular plexiform lesions involved dysregulated EC growth rather than abnormal proliferation of VSMC [52]. Further, an EC-reporter mouse with pulmonary hypertension showed that the immunostaining for α-SMA was colocalized with a small percentage of GFP-labeled cells of EC lineage, suggesting that cells from pulmonary vascular endothelial origin could undergo EndMT and lead to neointimal formation [53]. In addition, endothelial Klf-4 and Oct-4 are evident regulators of endothelial homeostasis, as EC knockout mice for these transcription factors show EndMT, with oxidative stress and inflammatory markers, rendering them more susceptible to neointima formation and plaque lesion instability [16,37,38]. We have previously shown that male BPH/2J mice at an early age (6-week-old) present with low Klf-4 levels in MRA before the rise of their systolic blood pressures [13], suggesting that dysregulation of endogenous vascular Klf-4 could contribute to the ECs phenotype transitioning in BPH/2J arteries.

Other recognized EndMT activators are oxidative stress and ROS. Studies have demonstrated that in primary human umbilical vein ECs (HUVECs) exposed to hydrogen peroxide (H_2_O_2_), the EndMT is observed and is mediated by TGF-β and Smad3, p38, and nuclear factor-kappa B (NF-κB) activation [54]. In addition, a transgenic mouse model with endothelial-specific Nox2 overexpression has been shown to have increased EndMT compared to wild-type mice [55]. OSK treatment of human hypertensive ECs was effective in decreasing ROS generation and improving NO production and bioavailability, suggesting that part of the beneficial effects induced by OSK *in vivo* involves the regulation of ROS production in arteries from BPH/2J mice, thereby also preventing EndMT.

Initial studies of EndMT suggested that it was a permanent process of cellular transdifferentiation into mesenchymal phenotype. However, new findings have demonstrated that cells undergoing EndMT pass through intermediate stages referred to as partial EndMT [21]. In these circumstances, cardiac fibroblasts undergo mesenchymal-to-endothelial-transition (MEndoT) to generate *de novo* ECs in the damaged heart and demonstrate that MEndoT can be augmented to enhance cardiac repair through neovascularization [56]. In 2015, Muir and co-workers [57] proposed that overexpressing Klf-4 in islet-enriched pancreatic cells to induce cell rejuvenation could represent an innovative strategy in regenerative medicine for type 1 diabetes. To this end, the authors observed that Klf-4 induced morphological changes, down-regulation of mesenchymal markers, and re-expression of both epithelial and pancreatic cell markers including insulin and transcription factors to β-cells. However, these effects were transient mainly due to increased apoptosis. Our OSK approach allowed us to obtain ECs with progenitor characteristics that prevented EndMT without inducing cellular death. *In vivo* treatment caused a reduction in mouse blood pressure values 10 days post-treatment, suggesting a more permanent effect after the overexpression of the three transcription factors together. To the best of our knowledge, no other specific EC reprogramming through the OSK strategy or the *in vivo* recovery of ECs from EndMT has been reported in hypertension, and this represents an innovative treatment for hypertension. These results provide the first direct evidence that OSK can promote reprogramming of ECs *in vivo* and it plays a protective role by preventing the deleterious effects induced by *in vivo* EndMT and vascular remodeling elicited by hypertension. This innovative therapeutic strategy may shed new light on the field of regenerative medicine in cardiovascular diseases.

## Novelty and Significance

### What is known?

- Hypertension affects 46.7% of adult U.S. population, and small artery remodeling and endothelial dysfunction are the major pathophysiological characteristic of this cardiovascular disease;
- Growing evidence supports a robust and likely causal association between cardiovascular diseases and the presence of EndMT;
- EndMT is a cellular transdifferentiation process in ECs which partially lose their identity and acquire additional mesenchymal phenotypes, including the gain of contractile properties.

### What new information does this article contribute?

- We showed that single-short partial reprogramming of ECs, via overexpression of OSK transcription factors, was able to bring ECs back to a youthful phenotype in hypertension;
- OSK treatment of hypertensive mice normalized blood pressure and resistance arteries hypercontractility, via the attenuation of EndMT and elastin breaks;
- OSK-treated human ECs from hypertensive patients showed high eNOS activation with high NO production, and the single-cell RNA analysis showed that OSK alleviated EC senescence and EndMT, restoring their phenotypes.

To the best of our knowledge, no other specific EC reprogramming through the OSK strategy or the *in vivo* recovery of ECs from EndMT has been reported in hypertension, and this represents an innovative treatment for hypertension. These results provide the first direct evidence that OSK can promote reprogramming of ECs *in vivo*. It plays a protective role by preventing the deleterious effects induced by *in vivo* EndMT and vascular remodeling elicited by hypertension. Overall, these data indicate that OSK treatment and EC reprogramming can decrease blood pressure and reverse hypertension-induced vascular damage. This innovative therapeutic strategy may shed new light on the field of regenerative medicine in cardiovascular diseases.

## Supporting information

Unedited WB membranes

## Acknowledgements

The authors thank the Instrumental Research Facility (IRF) at the School of Medicine, University of South Carolina. The research reported in this publication was supported by the National Institute of General Medical Sciences of the National Institutes of Health under Award Number P20GM103499; by the Office of The Director of the National Institutes of Health under Award Number 1S10OD032271-01 to J.L.K., and by the Columbia VA Healthcare System. The content is solely the responsibility of the authors and does not necessarily represent the official views of the National Institutes of Health (RRID:SCR_024955). We are also grateful for Dr. Sujit Pujhari and the Viral Vector Core, at the School of Medicine, University of South Carolina for lentiviral vector preparation, and Dr. Michael Shtutman and USC CTT COBRE Functional Genomics Core, at the College of Pharmacy, University of South Carolina for assistance with RNA sequencing data. All graphical abstracts were generated using BioRender.com.

## Sources of Funding

We gratefully acknowledge funding support from the National Institutes of Health (NIH) (R00GM118885 and R01HL149762 to CFW) and Alzheimer’s Association (AARG-NTF-23-1145090 to CFW); R00HL151889 to CGM; R01AR073172, 1R21AR083066 (NIAMS) and HT9425-23-1-008 (DoD) to WT.

## Disclosures

The authors declare no competing interests.

### Supplemental Material

Supplemental Methods

Table S1

Figure S1 – S5

References 25; 58 – 64

**Correspondence and requests for materials** should be addressed to Camilla F. Wenceslau.

## Notes

### Competing Interest Statement

The authors have declared no competing interest.

## References

[1] Park, J. B., Schiffrin, E. L. Small artery remodeling is the most prevalent (earliest?) form of target organ damage in mild essential hypertension. Hypertens. 19 (5): 921–930 (2001). Doi: 10.1097/00004872-200105000-00013.

[2] Mulvany, M. J. Small artery remodelling in hypertension: causes, consequences and therapeutic implications. Med. Biol. Eng. Comput. 46 (5): 461–467 (2008). Doi: 10.1007/s11517-008-0305-3.

[3] Ranchoux, B., Antigny, F., Rucker-Martin, C., Hautefort, A., Péchoux, C., Bogaard, H. J., Dorfmüller, P., Remy, S., Lecerf, F., Planté, S., Chat, S., Fadel, E., Houssaini, A., Anegon, I., Adnot, S., Simonneau, G., Humbert, M., Cohen-Kaminsky, S., Perros, F. Endothelial-to-mesenchymal transition in pulmonary hypertension. Circulation 131 (11): 1006–1018 (2015). Doi: 10.1161/CIRCULATIONAHA.114.008750.

[4] Tombor, L. S., John, D., Glaser, S. F., Luxán, G., Forte, E., Furtado, M., Rosenthal, N., Baumgarten, N., Schulz, M. H., Wittig, J., Rogg, E. M., Manavski, Y., Fischer, A., Muhly-Reinholz, M., Klee, K., Looso, M., Selignow, C., Acker, T., Bibli, S. I., Fleming, I., Patrick, R., Harvey, R. P., Abplanalp, W. T., Dimmeler, S. Single cell sequencing reveals endothelial plasticity with transient mesenchymal activation after myocardial infarction. Nat. Commun. 12 (1): 681 (2021). Doi: 10.1038/s41467-021-20905-1.

[5] Wang, E., Chen, S., Wang, H., Chen, T., Chakrabarti, S. Non-coding RNA-mediated endothelial-to-mesenchymal transition in human diabetic cardiomyopathy, potential regulation by DNA methylation. Cardiovasc. Diabetol. 22 (1): 303 (2023). Doi: 10.1186/s12933-023-02039-4.

[6] Zhao, J., Zhao, C., Yang, F., Jiang, Z., Zhu, J., Yao, W., Pang, W., Zhou, J. DNMT1 mediates the disturbed flow-induced endothelial to mesenchymal transition through disrupting β-alanine and carnosine homeostasis. Theranostics 13 (13): 4392–4411 (2023). Doi: 10.7150/thno.84427.

[7] Zhu, X., Wang, Y., Soaita, I., Lee, H. W., Bae, H., Boutagy, N., Bostwick, A., Zhang, R. M., Bowman, C., Xu, Y., Trefely, S., Chen, Y., Qin, L., Sessa, W., Tellides, G., Jang, C., Snyder, N. W., Yu, L., Arany, Z., Simons, M. Acetate controls endothelial-to-mesenchymal transition. Cell Metab. 35 (7): 1163–1178.e10 (2023). Doi: 10.1016/j.cmet.2023.05.010.

[8] Cooley, L. S., Edwards, D. R. New insights into the plasticity of the endothelial phenotype. Biochem. Soc. Trans. 39 (6): 1639–1643 (2011). Doi: 10.1042/BST20110723

[9] Dejana, E., Hirschi, K. K., Simons, M. The molecular basis of endothelial cell plasticity. Nat. Commun. 8: 14361 (2017). Doi: 10.1038/ncomms14361.

[10] Lin, A., Brittan, M., Baker, A. H., Dimmeler, S., Fisher, E. A., Sluimer, J. C., Misra, A. Clonal expansion in cardiovascular pathology. JACC Basic Transl. Sci. 9 (1): 120–144 (2023). Doi: 10.1016/j.jacbts.2023.04.008.

[11] Takahashi, K., Yamanaka, S. Induction of pluripotent stem cells from mouse embryonic and adult fibroblast cultures by defined factors. Cell 126(4): 663–676 (2006). Doi: 10.1016/j.cell.2006.07.024.

[12] Lu, Y., Brommer, B., Tian, X., Krishnan, A., Meer, M., Wang, C., Vera, D. L., Zeng, Q., Yu, D., Bonkowski, M. S., Yang, J. H., Zhou, S., Hoffmann, E. M., Karg, M. M., Schultz, M. B., Kane, A. E., Davidsohn, N., Korobkina, E., Chwalek, K., Rajman, L. A., Church, G. M., Hochedlinger, K., Gladyshev, V. N., Horvath, S., Levine, M. E., Gregory-Ksander, M. S., Ksander, B. R., He, Z., Sinclair, D. A. Reprogramming to recover youthful epigenetic information and restore vision. Nature 588 (7836): 124–129 (2020). Doi: 10.1038/s41586-020-2975-4.

[13] Pernomian, L., Tan, W., McCarthy, C., Wenceslau, C.F. Reprogramming endothelial and vascular smooth muscle cells to prevent and treat hypertension. Med. Hypotheses 179: 111162 (2023). Doi: 10.1016/j.mehy.2023.111162.

[14] Madden, S. K., Araujo, A. D., Gerhardt, M., Fairlie, D. P., Mason, J. M. Taking the Myc out of cancer: toward therapeutic strategies to directly inhibit c-Myc. Mol. Cancer 20: 3 (2021). Doi: 10.1186/s12943-020-01291-6.

[15] Dull, T., Zufferey, R., Kelly, M., Mandel, R. J., Nguyen, M., Trono, D., Naldini, L. A third-generation lentivirus vector with a conditional packaging system. J. Virol. 72 (11): 8463–8471 (1998). Doi: 10.1128/JVI.72.11.8463-8471.1998.

[16] Mastej, V., Axen, C., Wary, A., Minshall, R. D., Wary, K. K. A requirement for Krüppel like factor-4 in the maintenance of endothelial cell quiescence. Front. Cell. Dev. Biol. 10: 1003028 (2022). Doi: 10.3389/fcell.2022.1003028.

[17] Pinto, D. M. S., Flaus, A. Structure and function of histone H2AX. Subcell. Biochem. 50: 55–78 (2010). Doi: 10.1007/978-90-481-3471-7_4.

[18] Peichev, M., Naiyer, A. J., Pereira, D., Zhu, Z., Lane, W. J., Williams, M., Oz, M. C., Hicklin, D. J., Witte, L., Moore, M. A., Rafii, S. Expression of VEGFR-2 and AC133 by circulating human CD34(+) cells identifies a population of functional endothelial precursors. Blood 95 (3): 952–958 (2000). Doi: 10.1182/blood.V95.3.952.003k27_952_958.

[19] Shmelkov, S. V., St. Clair, R., Lyden, D., Rafii, S. AC133/CD133/Prominin-1. Int. J. Biochem. Cell. Biol. 37 (4): 715–719 (2005). Doi: 10.1016/j.biocel.2004.08.010.

[20] Friedrich EB, Walenta K, Scharlau J, Nickenig G, Werner N. CD34^-^/CD133^+^/VEGFR-2^+^ endothelial progenitor cell subpopulation with potent vasoregenerative capacities. Circ. Res. 17; 98 (3): e20–25 (2006). Doi: 10.1161/01.RES.0000205765.28940.93.

[21] Piera-Velazquez, S., Jimenez, S. A. Endothelial to mesenchymal transition: role in physiology and in the pathogenesis of human diseases. Physiol. Rev. 99 (2): 1281–1324 (2019). Doi: 10.1152/physrev.00021.2018.

[22] Kovacic, J. C., Dimmeler, S., Harvey, R. P., Finkel, T., Aikawa, E., Krenning, G., Baker, A. H. Endothelial to mesenchymal transition in cardiovascular disease: JACC state-of-the-art review. J. Am. Coll. Cardiol. 73 (2): 190–209 (2019). Doi: 10.1016/j.jacc.2018.09.089.

[23] Liang, G., Wang, S., Shao, J., Jin, Y. J., Xu, L., Yan, Y., Günther, S., Wang, L., Offermanns, S. Tenascin-X mediates flow-induced suppression of EndMT and atherosclerosis. Circ. Res. 130 (11): 1647–1659 (2022). Doi: 10.1161/CIRCRESAHA.121.320694.

[24] Adjuto-Saccone, M., Soubeyran, P., Garcia, J., Audebert, S., Camoin, L., Rubis, M., Roques, J., Binétruy, B., Iovanna, J. L., Tournaire, R. TNF-α induces endothelial-mesenchymal transition promoting stromal development of pancreatic adenocarcinoma. Cell. Death Dis. 12 (7): 649 (2021). Doi: 10.1038/s41419-021-03920-4.

[25] Schlager, G. Selection for blood pressure levels in mice. Genetics 76 (3), 537–549 (1974). Doi: 10.1093/genetics/76.3.537.

[26] Nguyen, V., Gao, C., Hochman, M. L., Kravitz, J., Chen, E. H., Friedman, H. I., Wenceslau, C. F., Chen, D., Wang, Y., Nelson, J. S., Jegga, A. G., Tan, W. Endothelial cells differentiated from patient dermal fibroblast-derived induced pluripotent stem cells resemble vascular malformations of port-wine birthmark. Br. J. Dermatol. 189 (6): 780–783 (2023). Doi: 10.1093/bjd/ljad330.

[27] Klisic, A., Kavaric, N., Vujcic, S., Spasojevic-Kalimanovska, V., Ninic, A., Kotur-Stevuljevic, J. Endocan and advanced oxidation protein products in adult population with hypertension. Eur. Rev. Med. Pharmacol. Sci. 24 (12): 7131–7137 (2020). Doi: 10.26355/eurrev_202006_21707.

[28] Xiao, X., Jiang, H., Wei, H., Zhou, Y., Ji, X., Zhou, C. Endothelial senescence in neurological diseases. Aging Dis. 14 (6): 2153–2166 (2023). Doi: 10.14336/AD.2023.0226-1.

[29] Struewing, I. T., Durham, S. N., Barnett, C. D., Mao, C. D. Enhanced endothelial cell senescence by lithium-induced matrix metalloproteinase-1 expression. J. Biol. Chem. 284 (26): 17595–17606 (2009). Doi: 10.1074/jbc.M109.001735.

[30] Matthaei, M., Meng, H., Meeker, A. K., Eberhart, C. G., Jun, A. S. Endothelial Cdkn1a (p21) overexpression and accelerated senescence in a mouse model of Fuchs endothelial corneal dystrophy. Invest. Ophthalmol. Vis. Sci. 53 (10): 6718–6727 (2012). Doi: 10.1167/iovs.12-9669.

[31] Sanada, F., Taniyama, Y., Muratsu, J., Otsu, R., Shimizu, H., Rakugi, H., Morishita, R. IGF binding protein-5 induces cell senescence. Front. Endocrinol. (Lausanne) 9: 53 (2018). Doi: 10.3389/fendo.2018.00053.

[32] Sun, X., Feinberg, M. W. Vascular endothelial senescence: pathobiological insights, emerging long noncoding RNA targets, challenges and therapeutic opportunities. Front. Physiol. 12: 693067 (2021). Doi: 10.3389/fphys.2021.693067.

[33] Whelton, P. K., Carey, R. M., Aronow, W. S., Casey Jr., D. E., Collins, K. J., Himmelfarb, C. D., DePalma, S. M., Gidding, S., Jamerson, K. A., Jones, D. W., MacLaughlin, E. J., Muntner, P., Ovbiagele, B., Smith Jr., S. C., Spencer, C. C., Stafford, R. S., Taler, S. J., Thomas, R. J., Williams Sr., K. A., Williamson, J. D., Wright Jr., J. T. 2017 ACC/AHA/AAPA/ABC/ACPM/AGS/APhA/ASH/ASPC/NMA/PCNA Guideline for the prevention, detection, evaluation, and management of high blood pressure in adults: a report of the American College of Cardiology/American Heart Association task force on clinical practice guidelines. Hypertension 71 (6): e13–e115 (2018). Doi: 10.1161/HYP.0000000000000065.

[34] Martin, S. S., Aday, A. W., Almarzooq, Z. I., Anderson, C. A. M., Arora, P., Avery, C. L., Baker-Smith, C. M., Barone Gibbs, B., Beaton, A. Z., Boehme, A. K., Commodore-Mensah, Y., Currie, M. E., Elkind, M. S. V., Evenson, K. R., Generoso, G., Heard, D. G., Hiremath, S., Johansen, M. C., Kalani, R., Kazi, D. S., Ko, D., Liu, J., Magnani, J. W., Michos, E. D., Mussolino, M. E., Navaneethan, S. D., Parikh, N. I., Perman, S. M., Poudel, R., Rezk-Hanna, M., Roth, G. A., Shah, N. S., St-Onge, M. P., Thacker, E. L., Tsao, C. W., Urbut, S. M., Van Spall, H. G. C., Voeks, J. H., Wang, N. Y., Wong, N. D., Wong, S. S., Yaffe, K., Palaniappan, L. P.; American Heart Association Council on Epidemiology and Prevention Statistics Committee and Stroke Statistics Subcommittee. 2024 Heart Disease and Stroke Statistics: A Report of US and Global Data From the American Heart Association. Circulation 149 (8): e347-e913 (2024). Doi: 10.1161/CIR.0000000000001209.

[35] Diaz Brinton, R. Minireview: translational animal models of human menopause: challenges and emerging opportunities. Endocrinology 153 (8): 3571–3578 (2012). Doi: 10.1210/en.2012-1340.

[36] Cagnacci, A., Venier, M. The controversial history of hormone replacement therapy. Medicina (Kaunas) 55 (9): 602 (2019). Doi: 10.3390/medicina55090602.

[37] Zhou, G., Hamik, A., Nayak, L., Tian, H., Shi, H., Lu, Y., Sharma, N., Liao, X., Hale, A., Boerboom, L., Feaver, R. E., Gao, H., Desai, A., Schmaier, A., Gerson, S. L., Wang, Y., Atkins, G. B., Blackman, B. R., Simon, D. I., Jain, M. K. Endothelial Kruppel-like factor 4 protects against atherothrombosis in mice. J. Clin. Invest. 122 (12): 4727–4731 (2012). Doi: 10.1172/JCI66056.

[38] Shin, J., Tkachenko, S., Chaklader, M., Pletz, C., Singh, K., Bulut, G. B., Han, Y. M., Mitchell, K., Baylis, R. A., Kuzmin, A. A., Hu, B., Lathia, J. D., Stenina-Adognravi, O., Podrez, E., Byzova, T. V., Owens, G. K., Cherepanova, O. A. Endothelial OCT4 is atheroprotective by preventing metabolic and phenotypic dysfunction. Cardiovasc. Res. 118 (11): 2458–2477 (2022). Doi: 10.1093/cvr/cvac036.

[39] Zhu, Y. T., Li, F., Han, B., Tighe, S., Zhang, S., Chen, S. Y., Liu, X., Tseng, S. C. Activation of RhoA-ROCK-BMP signaling reprograms adult human corneal endothelial cells. J. Cell. Biol. 206: 799–811 (2014). Doi: 10.1083/jcb.201404032.

[40] Yao, J., Guihard, P. J., Blazquez-Medela, A. M., Guo, Y., Liu, T., Bostrom, K. I., Yao, Y. Matrix Gla protein regulates differentiation of endothelial cells derived from mouse embryonic stem cells. Angiogenesis 19:1–7 (2016). Doi: 10.1007/s10456-015-9484-3.

[41] Moonen, J. R., Chappell, J., Shi, M., Shinohara, T., Li, D., Mumbach, M. R., Zhang, F., Nair, R. V., Nasser, J., Mai, D. H., Taylor, S., Wang, L., Metzger, R. J., Chang, H. Y., Engreitz, J. M., Snyder, M. P., Rabinovitch, M. KLF4 recruits SWI/SNF to increase chromatin accessibility and reprogram the endothelial enhancer landscape under laminar shear stress. Nat. Commun. 13 (1): 4941 (2022). Doi: 10.1038/s41467-022-32566-9.

[42] Camerini-Otero, R. D. and Felsenfeld, G. Histone H3 disulfide dimers and nucleosome structure. Proc. Natl. Acad. Sci. U.S.A. 74 (12), 5519–5523 (1977). Doi: 10.1073/pnas.74.12.5519.

[43] Prado, F., Jimeno-González, S., Reyes, J. C. Histone availability as a strategy to control gene expression. RNA Biol. 14 (3): 281–286 (2017). Doi: 10.1080/15476286.2016.1189071.

[44] Chen, K., Long, Q., Xing, G., Wang, T., Wu, Y., Li, L., Qi, J., Zhou, Y., Ma, B., Schöler, H. R., Nie, J., Pei, D., Liu, X. Heterochromatin loosening by the Oct4 linker region facilitates Klf4 binding and iPSC reprogramming. EMBO J. 39 (1): e99165 (2020). Doi: 10.15252/embj.201899165.

[45] Chen, J., Chen, X., Li, M., Liu, X., Gao, Y., Kou, X., Zhao, Y., Zheng, W., Zhang, X., Huo, Y., Chen, C., Wu, Y., Wang, H., Jiang, C., Gao, S. Hierarchical Oct4 binding in concert with primed epigenetic rearrangements during somatic cell reprogramming. Cell Rep. 14 (6): 1540–1554 (2016). Doi: 10.1016/j.celrep.2016.01.013.

[46] Li, D., Shu, X., Zhu, P., Pei, D. Chromatin accessibility dynamics during cell fate reprogramming. EMBO Rep. 22 (2): e51644 (2021). Doi: 10.15252/embr.202051644.

[47] Malik, V., Glaser, L. V., Zimmer, D., Velychko, S., Weng, M., Holzner, M., Arend, M., Chen, Y., Srivastava, Y., Veerapandian, V., Shah, Z., Esteban, M. A., Wang, H., Chen, J., Schöler, H. R., Hutchins, A. P., Meijsing, S. H., Pott, S., Jauch, R. Pluripotency reprogramming by competent and incompetent POU factors uncovers temporal dependency for Oct4 and Sox2. Nat. Commun. 10 (1): 3477 (2019). Doi: 10.1038/s41467-019-11054-7.

[48] Redon, C., Pilch, D., Rogakou, E., Sedelnikova, O., Newrock, K., Bonner, W. Histone H2A variants H2AX and H2AZ. Curr. Opin. Genet. Dev. 12 (2): 162–169 (2002). Doi: 10.1016/s0959-437x(02)00282-4.

[49] Graves, R. A., Marzluff, W. F., Giebelhaus, D. H., Schultz, G. A. Quantitative and qualitative changes in histone gene expression during early mouse embryo development. Proc. Natl. Acad. Sci. U.S.A. 82 (17): 5685–5689 (1985). Doi: 10.1073/pnas.82.17.5685.

[50] Karnavas, T., Pintonello, L., Agresti, A., Bianchi, M. E. Histone content increases in differentiating embryonic stem cells. Front. Physiol. 5: 330 (2014). Doi: 10.3389/fphys.2014.00330

[51] Yang, C., Zhang, Z. H., Li, Z. J., Yang, R. C., Qian, G. Q., Han, Z. C. Enhancement of neovascularization with cord blood CD133^+^ cell-derived endothelial progenitor cell transplantation. Thromb. Haemost. 91 (6): 1202–1212 (2004). Doi: 10.1160/TH03-06-0378.

[52] Tuder, R. M., Groves, B., Badesch, D. B., Voelkel, N. F. Exuberant endothelial cell growth and elements of inflammation are present in plexiform lesions of pulmonary hypertension. Am. J. Pathol. 144 (2): 275–285 (1994). PMID: 7508683.

[53] Qiao, L., Nishimura, T., Shi, L., Sessions, D., Thrasher, A., Trudell, J. R., Berry, G. J,, Pearl, R. G., Kao, P. N. Endothelial fate mapping in mice with pulmonary hypertension. Circulation 129 (6): 692–703 (2014). Doi: 10.1161/CIRCULATIONAHA.113.003734.

[54] Montorfano, I., Becerra, A., Cerro, R., Echeverría, C., Sáez, E., Morales, M. G., Fernández, R., Cabello-Verrugio, C., Simon F. Oxidative stress mediates the conversion of endothelial cells into myofibroblasts via a TGF-β1 and TGF-β2-dependent pathway. Lab. Invest. 94 (10): 1068–1082 (2014). Doi: 10.1038/labinvest.2014.100.

[55] Murdoch, C. E., Chaubey, S., Zeng, L., Yu, B., Ivetic, A., Walker, S. J., Vanhoutte, D., Heymans, S., Grieve, D. J., Cave, A. C., Brewer, A. C., Zhang, M., Shah, A. M. Endothelial NADPH oxidase-2 promotes interstitial cardiac fibrosis and diastolic dysfunction through proinflammatory effects and endothelial-mesenchymal transition. J. Am. Coll. Cardiol. 63 (24): 2734–2741 (2014). Doi: 10.1016/j.jacc.2014.02.572.

[56] Ubil, E., Duan, J., Pillai, I. C., Rosa-Garrido, M., Wu, Y., Bargiacchi, F., Lu, Y., Stanbouly, S., Huang, J., Rojas, M., Vondriska, T. M., Stefani, E., Deb, A. Mesenchymal-endothelial transition contributes to cardiac neovascularization. Nature 514 (7524): 585–590 (2014). Doi: 10.1038/nature13839.

[57] Muir, K. R., Lima, M. J., Docherty, H. M., McGowan, N. W., Forbes, S., Heremans, Y., Forbes, S. J., Heimberg, H., Casey, J., Docherty, K. Krüppel-like factor 4 overexpression initiates a mesenchymal-to-epithelial transition and redifferentiation of human pancreatic cells following expansion in long term adherent culture. PLoS One 10 (10): e0140352 (2015). Doi: 10.1371/journal.pone.0140352.

[58] Uddin, M. H., Al-Hallak, M. N., Khan, H. Y., Aboukameel, A., Li, Y., Bannoura, S. F., Dyson, G., Kim, S., Mzannar, Y., Azar, I., Odisho, T., Mohamed, A., Landesman, Y., Kim, S., Beydoun, R., Mohammad, R. M., Philip, P. A., Shields, A. F., Azmi, A. S. Molecular analysis of XPO1 inhibitor and gemcitabine-nab-paclitaxel combination in KPC pancreatic cancer mouse model. Clin. Transl. Med. 13 (12): e1513 (2023). Doi: 10.1002/ctm2.1513.

[59] Wolf, F. A., Angerer, P., Theis, F. J. SCANPY: large-scale single-cell gene expression data analysis. Genome Biol. 19 (1): 15 (2018). Doi: 10.1186/s13059-017-1382-0.

[60] Korsunsky, I., Millard, N., Fan, J., Slowikowski, K., Zhang, F., Wei, K., Baglaenko, Y., Brenner, M., Loh, P. R., Raychaudhuri, S. Fast, sensitive and accurate integration of single-cell data with Harmony. Nat. Methods. 16 (12): 1289–1296 (2019). Doi: 10.1038/s41592-019-0619-0.

[61] Liao, Y., Smyth, G. K., Shi, W. Feature Counts: an efficient general-purpose program for assigning sequence reads to genomic features. Bioinformatics 30 (7): 923–930 (2014). Doi:10.1093/bioinformatics/btt656.

[62] Robinson, M. D., McCarthy, D. J., Smyth, G. K. EdgeR: a Bioconductor package for differential expression analysis of digital gene expression data. Bioinformatics 26 (1): 139–140 (2010). Doi:10.1093/bioinformatics/btp616.

[63] McCarthy, D. J., Chen, Y., Smyth, G. K. Differential expression analysis of multifactor RNA-Seq experiments with respect to biological variation. Nucleic Acids Res. 40 (10): 4288–4297 (2012). Doi:10.1093/nar/gks042.

[64] Benjamini, Y., and Hochberg, Y. Controlling the false discovery rate: A practical and powerful approach to multiple testing. Journal of the Royal Statistical Society: Series B (Methodological) 57: 289–300 (1995). Doi: 10.1111/j.2517-6161.1995.tb02031.x.

[65] Paxinos and Franklin’s The Mouse Brain in Stereotaxic Coordinates, Compact: *The Coronal Plates and Diagrams* 5^th^ Edition by Franklin MA PhD, Keith B.J.; Paxinos AO (BA MA PhD DSc) FASSA FAA, George. ISBN-10: 0128161590. ISBN-13: 9780128161593; Edition: 5; Released: Jun 18, 2019; Publisher: Academic.

[66] Biancardi, V. C., Campos, R. R., Stern, J. E. Altered balance of gamma-aminobutyric acidergic and glutamatergic afferent inputs in rostral ventrolateral medulla-projecting neurons in the paraventricular nucleus of the hypothalamus of renovascular hypertensive rats. J. Comp. Neurol. 518 (5): 567–585 (2010). Doi: 10.1002/cne.22256.

[67] Costa, T. J., Jiménez-Altayó, F., Echem, C., Akamine, E. H., Tostes, R., Vila, E., Dantas, A. P., Carvalho, M. H. C. Late onset of estrogen therapy impairs carotid function of senescent females in association with altered prostanoid balance and upregulation of the variant ERα36. Cells 8 (10): 1217 (2019). Doi: 10.3390/cells8101217.

