## Supplemental material for "A Single-Short Partial Reprogramming of the Endothelial Cells decreases Blood Pressure via attenuation of EndMT in Hypertensive Mice"

### Supplemental Methods

#### Lentiviral vector (LV) production

Human embryonic kidney (HEK293T) cells were cultured in MEM with 10% fetal bovine serum (FBS) in a 225 $cm^2$ flask at 37 °C with 5% $CO_2$ to generate lentivirus. For co-transfection, three packaging plasmids (psPAX2, pRSV-Rev, and pMD2.G) and lentiviral vector for mouse Cadherin 5-(also known as Cdh5 or VE-cadherin)-Oct4-Sox2-Klf4-EGFP (here, referred as LV-OSK) or the Cdh5-EGFP (referred as LVCO) were mixed with 200mM PEI in PBS and incubated for 10 min at room temperature. Subsequently, the mixture was added to HEK293T cells and incubated at 37°C with 5% CO2 overnight. The next day, the transfection medium was removed and replaced with a virus-production medium, MEM containing 2% FBS, 7.5% NaHCO3, 200 mM Glutamine, 100 mM Pyruvate, and 20% glucose. After 72 h post-transfection, the incubated supernatant was collected and centrifuged for 4 min at 4°C and 4000 rpm, filtered through a 0.45 μm filter, followed by high-speed centrifugation for 2 h at 4°C and 20,000 rpm. The virus pellet was suspended in 1 ml of virus suspension media and layered over 500 µl of 20% sucrose solution, followed by high-speed centrifugation for 2 h at 4°C and 20,000 rpm. The virus pellet was suspended in 1 ml of virus suspension media and laid over 500 µl of 20% sucrose solution, followed by high-speed centrifugation for 2 h at 4°C and 20,000 rpm. The final virus pellet was suspended in 100 µl of VSM, aliquoted, and stored at -80°C for further use. A standardized method was used to titrate the virus and expressed as transfection units per milliliter (TU/ml). LV production and utilization in cells and mice had the School of Medicine, University of South Carolina ethics committee approval (IACUC #2596-101690-041122, and IBC protocol # 300322).

**Mouse and Human EC culture**

C57Bl/6 mouse intestinal mesenteric primary endothelial cells (Accegen, ABC-TC3197) were used in the following experiments with phosphate-buffered saline (PBS), LVCO or LV-OSK. Briefly, mouse endothelial cells were seeded at a density of $1x10^5$, $4x10^5$ or $1x10^6$ cells/plate depending on the respective assay, using Endothelial Cell Medium (Cell Biologics, M1168 + supplement kit) with 5% fetal bovine serum (FBS) and 1% antibiotic-antimycotic solution, and were infected in suspension with LVCO or LV-OSK (stock solution at $2.81x10^8$ TU/mL) to obtain multiplicity of infection (MOI) 1. After 3 or 5 days, cells were washed with fresh endothelial cell medium, and collected for experiments.

The primary normotensive and hypertensive human aortic endothelial cell (HAoEC) were purchased from PromoCell. At 60% confluence, the HAoEC were infected with sendai virus (SeV) carried polycistronic Klf4-Oct3/4-Sox2 and Klf4 vectors (ThermoFisher, Waltham, MA, #A16517), here referred as OSK-SeV, to induce cell reprogram. The MOI of Klf4-Oct3/4-Sox2 and Klf4 was respectively 5:3. The

HAoEC was also infected with control SeV with EmGFP (Thermo Fisher Scientific, A16519), named as EGFP-SeV (control condition). Some experiments were performed with SeV with no EGFP to avoid green fluorescence interference. In this case, PBS was used as control condition, and OSK-SeV with no EGFP referred to the OSK factors overexpression in HAoEC. The ECs were collected at day 5 for further analysis.

**Animals**

All animal procedures and protocols used were approved by the Animal Care and Use Committee at the University of South Carolina School of Medicine. Experiments were conducted following the National Institutes of Health Guide for the Care and Use of Laboratory Animals and Animal Research Reporting of in Vivo Experiments (ARRIVE) guidelines. Male and female BPN/3J (RRID:IMSR_JAX:003004) and BPH/2J (RRID:IMSR_JAX:003005) mouse strains were obtained from The Jackson Laboratory and maintained as an inbred colony at the Animal Facility (School of Medicine Columbia, University of South Carolina). Male and female mice were used at 40-44 weeks of age for the intravenous treatment, and a female mice group at 30 weeks of age was used for the intraperitoneal treatment with the LVCO or LV-OSK. All mice were maintained on a 12-hour light/dark cycle with water and standard chow diet ad libitum.  Note that the sample size indicated per experiment is the number of independent mice used (experimental n). BPN/3J and BPH/2J were used as normotensive and hypertensive mice, respectively [25].

**Flow cytometry**

Treated mouse endothelial cells were analyzed for their EGFP positive ($EGFP^+$) signals, or for activation of apoptosis and necrosis, using the Pacific Blue Annexin V Apoptosis Detection Kit with 7-amino-actinomycin D (7-AAD) (BioLegend, #640926). Briefly, confluent treated-endothelial cells for 3 days with PBS, LVCO, or LV-OSK were collected with Trypsin-EDTA 0.25% (Gibco, #25200056) in polystyrene tubes for flow cytometry, and washed in Hanks' Balanced Salt Solution without calcium chloride, magnesium chloride and no phenol red (HBSS; Gibco, #14175-095) at 37°C, centrifuged at 1350 rpm for 5 min. The supernatant was discarded and for the $EGFP^+$ signals, cells were resuspended in fresh HBSS and analyzed using BD FACSymphony™ A5 Cell Analyzer (BD Biosciences) on blue laser (488 nm) and FITC detector (515/20). For the apoptosis and necrosis assay, $1x10^6$ cells/100 µL were resuspended in 100 µL annexin V binding buffer and 5 µL Pacific Blue Annexin V and 5 µL 7-AAD viability staining solution was added into tubes, gently vortexed, and incubated in the dark for 15 min at room temperature. After, 400 µL annexin V binding buffer was added in each tube, and cells were analyzed by flow cytometry using blue laser (488 nm) for 7-AAD detection at 670/30 nm long-pass filter, and violet laser (405 nm) for Pacific Blue detection at 450/50 nm. Blank cells were not incubated with Annexin V Pacific Blue or 7-AAD. DIVA Software was used for flow cytometry analysis.

**Wound healing assay**

PBS-, LVCO-, or LV-OSK-treated mouse endothelial cells (3 days) were plated on a 24-well plate with a fresh endothelial cell medium. In the next day, confluent wells were scratched on the bottom with a 10 µL sterile pipette tip. After, cells were placed in the cell incubator for 24 h at 37°C (5% CO2). Plates were photographed on a stereo microscope (Axiovert 5, Zeiss) to check for cellular growth (migration + proliferation) using Zen Software 3.5. Photos were taken after scratch and were referred as 0 h, and after 24 h. Images were analyzed by ImageJ Software to measure the % cell growth after 24 h compared to 0 h.

**Confocal microscopy and immunofluorescence**

Mouse ECs were treated with PBS, LVCO, or LV-OSK for 3 days (for CD31, CD133, CD34, and CD45 evaluation) or 5 days (for CD31, Oct-3/4, Ki67, Sox-2, and total histone H2Ax) prior to the immunofluorescence protocol. HAoEC were treated with PBS (control) or OSK-SeV (with no EGFP) for 5 days. The coronary arteries in the left ventricle from male BPN/3J or BPH/2J mice with no treatments (at 72 weeks of age), and MRA isolated from male and female BPN/3J or BPH/2J mice also with no treatments (at 72 weeks of age) were used to show the EndMT (CD31, α-SMA and DAPI staining). Brains (prefrontal cortex) isolated from i.v. treated LVCO or LV-OSK male and female mice were analyzed by their EGFP signals and CD31 expression. Briefly, one hemisphere of each animal was fixed in 4% paraformaldehyde (PFA) for 24 hours at 4°C. After fixation, the brains were cryoprotected at 4°C in 0.01 M PBS containing 15% sucrose followed by 30% sucrose for at least 72 hours. Serial coronal prefrontal cortex (between bregma 1.98 and 0.98 mm; [65]) slices of 30 μm were cut using a cryostat and collected in PBS 0.01M. The tissue was incubated with PBST (0.01 M PBS, 0.01% Triton X-100, 0.04% $NaN_3$), and 5% normal donkey serum followed by a primary antibody to stain endothelial cells (goat anti-CD31 polyclonal antibody, R&D systems, #AF3628; 1:200, overnight). Immunofluorescence was visualized by incubating the tissue with a secondary fluorescently labeled antibody for 4 hours (anti-goat Alexa-Fluor-594-conjugated donkey anti-goat, Jackson Immunoresearch, #705-585-147, 1:500). EGFP was readily detected using 488 nm excitation laser. Sections were washed and mounted with VECTASHIELD® Antifade Mounting Medium with DAPI (Vector Laboratories, USA). Negative control sections were run with the omission of the primary antibody, in which no fluorescence signal was observed. Full-thickness confocal z-stacks (1 μm interval, x20 objective) from three sections/mice were acquired with a Nikon Eclipse TE2000-E inverted microscope coupled to a Nikon A1 confocal laser. The optical sections were acquired with identical acquisition settings, and the two different lasers were emitted sequentially to avoid crosstalk. Immunoreactive signals of the maximum projection images were analyzed using ImageJ software. Background fluorescence intensity was obtained from the neuropil of each image, and a threshold was set to pass intensities approximately four times above the background signal. The density of the endothelial cells was calculated by dividing the overall CD31 intensity of fluorescence by the area occupied and expressed as a percentageConfocal microscopy images were acquired to analyze fluorescence intensity and the percent area of fluorescence intensity using a threshold paradigm through ImageJ [66].

In addition, thoracic aortas isolated from i.v. treated LVCO or LV-OSK male mice at 40-44 weeks of age (for CD31, α-SMA; DAPI and EGFP detection) were embedded in Tissue Tek® O.C.T. Compound (Sakura, #4583), frozen in liquid nitrogen, sectioned (5 μm) using Thermo Scientific Cryostat Microm HM525 (Thermo Scientific), and slides were analyzed by immunofluorescence. Briefly, cells were seeded on a 24-well plate with a glass coverslip in the bottom. For the Oct-3/4 and Ki67 assay, treated-confluent cells underwent the wound healing assay to stimulate proliferation and migration for 24h prior to immunofluorescence protocol. Samples were fixed with 4% paraformaldehyde (PFA) for 20 min, and permeabilized with 0.1% Triton X-100 and 0.01 M glycine for 1h at 37°C. Following, slides were incubated with 5% bovine serum albumin (BSA) for 15 min and subsequently incubated with 5% goat serum for 30 min at room temperature to reduce non-specific binding. After, samples were incubated overnight at 4°C with the following primary antibodies: goat anti-Oct-3/4 (1:100, Invitrogen, PA5-19307), rat anti-Sox-2 (1:100, Invitrogen, #14-9811-82), rabbit anti-Ki67 SP6 (1:100, Invitrogen, MA5-14520), rabbit anti-total histone H2Ax (1:100, Novus Biologicals, NB100-638), rat anti-CD31 Alexa Fluor 647 clone 390 (1:300, BioLegend, #102416), rabbit anti-CD133 (1:100, Novus Biologicals, NB120-16518), anti-CD34 MEC14.7 (1:100, Novus Biologicals, NB600-1071), rabbit anti-CD45 (1:100, Santa Cruz Technologies, sc-25590), or mouse anti-α-smooth muscle actin-DyLight 550 1A4-asm-1 (1:500, Novus Biologicals, NBP2-34522R). Secondary antibodies were used for 1 h at room temperature: goat anti-rabbit Alexa Fluor 647 (1:500, Invitrogen, A32733), goat anti-rat Alexa Fluor 647 (1:500, Invitrogen, A21247), donkey anti-goat Alexa Fluor 647 (1:500, Invitrogen, A32849), goat anti-rabbit Alexa Fluor 568 (1:500, Invitrogen, A11011), or goat anti-rabbit Texas red (1:300, Invitrogen, T2767), and samples were protected from light. Glass coverslips with cells or aortas were mounted with Fluoroshield™ with DAPI (Sigma Aldrich, F6057). Negative control represented cells or arteries with no primary antibody incubation. High-resolution images were acquired on confocal microscope (Stellaris 5 LIAchroic Confocal System, Leica), on 20x objective (for CD133 assay), or 63x objective. The LAS-X Software was used with the following configurations: 1024x1024, 400 Hz, 16 bits, *xyz* on sequential mode, and LAS-X or FIJI Software was used for image analysis. Colocalization analysis of CD31 (green) and α-SMA (red) in coronary arteries was performed on LAS-X Software, and % positive colocalization signal were shown in white in the images. Data presented as fluorescence intensity (A.U.) or % area were acquired on an area of 184.52 $\mu m^2$ (for 63x objective) or 581.25 $\mu m^2$ (for 20x objective) depending on the assay.

**Western blotting**

Mouse endothelial cells treated with PBS, LVCO, or LV-OSK for 3 days were evaluated by Western blotting. Additionally, 5 days post-infection EGFP-SeV (control) or EGFP-OSK-SeV, the normotensive and hypertensive HAoEC were serum-starved for 2 h before the experiment was performed. 50 ng/mL VEGF165 (Peprotech, Cranbury, NJ) was added to HAoEC for 30 mins. The cells were lysed with RIPA buffer for Western Blot analysis. Briefly, Klf-4, phosphorylated Serine 1177 (S1177) of endothelial nitric oxide synthase (eNOS), total eNOS and glutharaldehyde-phosphate dehydrogenase (GAPDH) protein expression were evaluated in mouse or human endothelial cells. Briefly, 50 µg protein of mouse endothelial cells treated with PBS, LVCO or LV-OSK, or 10 µg protein of HAoEC treated with EmGFP-SeV or EmGFP-OSK-SeV were loaded in 10-12% SDS-PAGE, and transferred to a nitrocellulose (for mouse endothelial cells) or PVDF membrane (for HAoEC), followed by blocking with 5% milk solution for 2h, and overnight incubation with the primary antibodies at 4° C: rabbit polyclonal anti-Klf4 (1:200; Abcam, ab129473; bands consist on monomeric form at 60-75 kDa, and dimeric form at ~100 kDa), rabbit polyclonal anti-total histone H3 (1:1000; Abcam, AB1791, observed bands between 15-20 kDa) or rabbit polyclonal anti-GAPDH (1:500; AB Clonal, #AC001, detected ~37 kDa) for mouse endothelial cells, or mouse monoclonal anti-phospho-(S1177)-eNOS (1:250, BD Biosciences, #612392; detected at ~140 kDa), and total rabbit polyclonal anti-total eNOS (1:250, Novus Biologicals, NB300-500; observed band ~140 kDa) for HaoEC. Secondary antibodies donkey anti-rabbit HRP or goat anti-mouse HRP (1:1,000; Jackson Immuno Research #711-035-152, and #115-035-062, respectively) were incubated for 1h at room temperature. Chemiluminescence was detected using Pierce™ ECL Western blotting substrate (Thermo Fisher Scientific, #32106). Ponceau red staining was used to check protein transfer after electrophoresis. GAPDH was used as loading control, and to normalize target bands. Data were analyzed by ImageJ Software (National Institutes of Health, NIH).

**Senescence-associated (SA) β-galactosidase assay**

Cellular senescence was evaluated in mouse endothelial cells by the SA β-galactosidase activity assay (Abcam, ab65351). Briefly, confluent mouse endothelial cells (passage 7) treated with PBS, LVCO, or LV-OSK for 120 h were plated on a 24-well plate with a fresh endothelial cell medium. In the next day, confluent cells were washed with PBS and fixed with 500 µL Fixative Solution III provided by the manufacturer, for 15 min at room temperature. After, cells were washed twice and 500 µL Staining Solution Mix was added in each well. The plate was covered and incubated overnight at 37°C. On the next day, cells were observed in a stereo microscope (20x objective, EVOS Cell Image Systems, Thermo Fisher Scientific) and blue-stained cells were considered senescent cells. Senescence was evaluated in basal levels. The negative control was represented by PBS-treated cells and fixed with the fixative solution without incubation with the specific staining solution. The number of senescent cells was counted using ImageJ Software.

**Digital spatial profiling (DSP) of aortas from LVCO and LV-OSK-treated mice at a transcript-proteomic scale**

DSP was performed on formalin-fixed paraffin-embedded (FFPE) thoracic aortas isolated from male mice, and samples were processed by City of Hope (California, CA, USA), as described by Uddin et al. [58]. Briefly, 5 µm aorta sections from the i.v. LVCO- or LV-OSK-treated (40-44-week-old) mice were affixed in the middle of a charged slide. The slide was baked at 60°C for 30 min and subjected to deparaffinization, rehydration, and antigen retrieval. To perform RNA profiling, a set of probes designed for mRNAs (Mouse Whole Transcriptome Atlas RNA v1.0) was applied to the sample and allowed to hybridize at 37°C overnight inside a hybridization oven. In preparation for protein profiling, subsequently, a tricolor fluorescence morphology marker panel was applied to the slides, targeting pan-cytokeratin (Pan-CK; 2 μg/mL; Novus Biologicals; NBP2-33200) to stain filamentous proteins of epithelial cells, CD45 (1:40 dilution; Novus Biologicals; NBP1-44763AF594) as a transmembrane protein of all differentiated hematopoietic cell marker, and SYTO83 (0.2 μM; ThermoFisher Scientific; S11364) for the nucleus. Specific oligo-conjugated primary antibodies cocktails were incubated with the slides. Every target probe was equipped with an exclusive photocleavable oligonucleotide barcode, which was amenable to quantification through next-generation sequencing. For both RNA and protein profiling, the slide was loaded into the profiler covered with an acquisition buffer. The utilization of multicolored morphology markers enabled the visualization of different compartments. This visualization guided the selection of regions of interest (ROIs; i.e., the endothelial or the vascular smooth muscle layers). After the one-step overnight incubation, the samples underwent 20x high-precision scanning using a GeoMx DSP system, followed by the selection of ROIs. The DNA oligonucleotides, which were attached to the profiling reagents, were released sequentially using ultraviolet illumination and gathered into individual wells on a 96-well plate. Subsequently, the collected DNA was subjected to Illumina library preparation. The expression levels were measured using an Illumina Sequencer and then analyzed using the DSP interactive software (GeomxTools, version 1.99.4; Advanced Genomics Core).

**RNA sequencing for human and mouse ECs**

For human ECs, it was used the ParseBioscience multiplexed scRNA-seq associated fastq files that contain reads from multiple samples due to an additional layer of multiplexing. Tracking of the sample origin is achieved using additional barcodes in R2. For this reason, the ParseBiosciences-Pipeline (v1.2.0) was used to align and quantify multiplexed reads into one cell by gene expression count matrix, using human genome GRCh38 as reference. The scRNA-seq data analysis was performed with the Scanpy (v1.9) package [59] in Python. Briefly, genes expressed in fewer than 100 cells were removed from further analysis. Cells expressing less than 100 genes or with a high (>0.15) mitochondrial genome transcript ratio were removed from further analysis. For downstream analysis, we used count per million normalizations (CPM) to control for library size difference in cells. After normalization, we used the ‘pp.highly_variable_genes’ command in Scanpy to find highly variable genes across all cells within each sample using default parameters. The data were then Z score normalized for each gene across all cells. Batch correction was performed using ‘harmonypy’ (https://github.com/slowkow/harmonypy), a python wrapper of the Harmony algorithm [60]. We then used the ‘pp.neighbors’ and the ‘tl.leiden (resolution = 0.5)’ command in Scanpy to partition the single cells into clusters. Differential expression analysis was performed using the Wilcoxon Rank-Sum test implemented in the ‘tl.rank_genes_groups’ function. Functional enrichment analysis and GSEA analysis were performed using the ‘GSEApy’ Python package (https://github.com/zqfang/GSEApy).

Mouse ECs were seeded at a density of 5x$10^5$ cells and treated for 24 h with TNF-α (100 nM) or vehicle (0.1 M phosphate buffer). After that, the cells were washed (0.1 M phosphate buffer) and subsequently centrifuged (300 xg for 5 min), the supernatant was removed and the cells were stored at −80°C. RNA sequencing RNA and library preparation, post-processing of the raw data, and data analysis were performed by the USC CTT COBRE Functional Genomics Core. RNAs were extracted with ZYMO Quick-RNA Microprep Kit R1050 (ZYMO, Irvine, CA, USA). RNA quality was validated on an RNA-1000 chip using Bioanalyzer (Agilent, Santa Clara, CA, USA). RNA libraries were prepared using an established protocol with the NEBNExt Ultra II Directional Library Prep Kit (NEB, Lynn, MA). Each library was made with one of the TruSeq barcode index sequences, and the sequencing was performed by Medgenome (Foster City, CA) on the Illumina NovaSeq platform (150 bp, pair-ended). Raw sequencing reads were aligned to the Mus Musculus genome GRCm38.p6 (GCA_000001635.8, ensemble release-102) using STAR v2.7.2b. The Samtools (v1.5) was used to convert aligned sam files to bam files and aligned reads were counted using the featureCounts command from the Subreads package [61]. Only reads mapped uniquely to the genome were used for gene expression analysis. Differential expression analysis was performed in R using the edgeR package [62]. Raw counts were normalized using the Trimmed Mean of M-values (TMM) method, and the normalized read counts were then fitted to a generalized linear model using the function glmFit [63]. Genewise tests for significant differential expression were performed using the function glmLRT. The P-value was then corrected for multiple testing using Benjamini-Hochberg's FDR [64].

**Nitric oxide (NO) and reactive oxygen species (ROS) production evaluation by DAF-FM/DA and DHE fluorescence**

In order to analyze the intracellular production of NO and ROS, HAoEC treated with PBS or OSK-SeV (with no EGFP) for 5 days, were incubated with the selective intracellular NO dye DAF-FM/DA (Invitrogen, D23844; 10 μM) or intracellular ROS dye Dihydroethidium (DHE, Invitrogen, D23107; 10 μM), respectively. Briefly, cells were washed with HBSS with calcium chloride, magnesium chloride, and no phenol red (Gibco; #14025092), and incubated at 37°C for 30 min in the dark with the appropriate dye. Slides were mounted and images were acquired on a confocal microscope (Stellaris 5 LIAchroic Confocal System, Leica; 20x objective) with LAS-X Software. For the NO assay, Acetylcholine (ACh, 1 μM, 5 min) was added before mounting the slides with Fluoromount™ (Sigma Aldrich, F4680), and images were acquired. ImageJ Software was used to quantify 30 different regions of interest (60 h x 48 w) and the intracellular fluorescence intensity of DAF-FM/DA or nuclear fluorescence intensity (A.U.) of DHE was evaluated. Negative controls represented cells with no DAF-FM/DA or DHE incubation.

**LVCO and LV-OSK treatment in mice**

One intravenous (i.v.) injection of LV carrying control plasmid (LVCO) or LV containing the three mouse transcription factors Oct-3/4, Sox-2 and Klf-4 (LV-OSK), both with cadherin 5 (Cdh5) and enhanced green fluorescent protein (EGFP) reporter, was performed in male and female BPN/3J and BPH/2J (40-44-week-old) mice through the tail vein. Briefly, mice were anaesthetized with 2% isoflurane and placed on a heating pad (37° C) platform. The tail vein was punctured and 100 μL (5 μL LVCO or LV-OSK + 95 μL sterile saline) was injected using a 30-gauge syringe. One separate experiment involved one intraperitoneal (i.p.) injection of 100 μL (5 μL LVCO or LV-OSK + 95 μL sterile saline) only in female BPN/3J and BPH/2J mice at 30 weeks of age. Regardless of the route of administration, mice were placed in their respective cages and monitored until recovered, and experiments were performed 10 days post-treatment.

**Echocardiography in mice**

Male and female BPH/2J and BPN/3J mice, i.v. treated with LVCO or LV-OSK and at 40-44 weeks of age were lightly anesthetized with 3% isoflurane (1 L/min 100% oxygen), using an anesthesia system that includes active scavenging, and maintained at ambient body temperature on the top of a heated stage. Once the animal lost its righting reflex, it was laid supine on a heated platform with its nose enveloped in a nosecone to keep the mouse anesthetized by 1.5% isoflurane. Heart rate was determined from a surface electrocardiogram. This regimen maintains heart rate at 400-500 bpm and the mice remain normothermic. This procedure ensured that in vivo measurements of left ventricular geometry and function were determined at physiologically relevant heart rates. Hair on the chest of the animal was removed, and covered with a 1 cm coating of gel solution to facilitate ultrasound transmission. Transthoracic echocardiography images were obtained by placement of an echocardiograph probe in the gel and moving throughout the thoracic region. From a transthoracic approach, 2-dimensional targeted M-mode echocardiographic recordings were obtained in the parasternal short and long axis views. Two-dimensional M-mode echocardiographic recordings were obtained using a 40 MHz scanning head with a spatial resolution of 30 μm on Visual Sonics VEVO 3100 High Resolution In Vivo Imaging System. Echocardiographic measurements were made as an average of at least five beats for each measurement. Left ventricle dimension and wall thickness were made at end-systole and end-diastole using the American Society of Echocardiography and established criteria for mice. Mean wall thickness was calculated as the average of the end-diastolic left ventricular anterior wall thickness (LVAW d) and left ventricular posterior wall thickness (LVPW d). Left ventricle mass was calculated using a standard formula on VEVO Lab Software. Left ventricle end-diastolic volume and end-systolic volume were determined using Simpson's method of disks and used to compute ejection fraction (EF). The pulse wave velocity (PWV) was calculated by placing the Doppler sample volume at the aortic arch using the total distance of the aortic arch, and the ascending and descending aortic flow velocities related to the corresponding QRS complex wave (transit time 1 and 2), and are showed as values in cm/s.

**Blood pressure measurement by left carotid catheterization**

Treated male and female BPN/3J or BPH/2J mice were anesthetized with 3% isofluorane (1 L/min 100% oxygen) and placed in the supine position on a warm pad. A small incision on the skin was made on the left side of the neck, near the mouse trachea, and the left carotid artery was exposed and carefully separated from other neighboring structures. A silk suture was placed distally for the complete ligation of the vessel. A second silk suture was placed proximally to allow temporary obstruction of blood flow. Finally, a third silk suture was placed loosely between the first two ligatures and a small incision (arteriotomy) was made between the first and the third suture. A tip of a catheter that had been pre-filled with 2% heparinized sterile saline was inserted into the carotid artery via the arteriotomy in the direction of the heart and secured in place by tying the third suture once the catheter had been advanced past the ligature.

The isoflurane anesthesia was reduced by 1%, and the pulsatile arterial pressure (PAP, in mmHg) was acquired for 20 minutes after the stabilization of the signal, using the LabChart 7 Software. After blood pressure measurements, blood was collected through the arterial catheter in chilled heparinized tubes under anesthesia (5% isoflurane). Plasma was obtained after centrifugation at 1,000 x g for 15 min at 4°C, and stored at -80°C until experiments. The systolic blood pressure (SBP, in mmHg) was calculated and analyzed using the LabChart 7 Software formulas. The SBP was analyzed as the mean of 10 different stable maximum values taken in 1-minute intervals. Data presented on Fig. 5 referred to i.v. treated male and female mice (40-44-week-old), and Extended Data Fig. 2 shows blood pressure values from i.p. treated female (30-week-old) mice.

**Vascular function in mesenteric resistance arteries (MRA)**

Second-order MRA (with an inner diameter up to 250 μm) were isolated from male and female LVCO- or LV-OSK-treated BPN/3J and BPH/2J mice (i.v.), and 2 mm length segments were mounted on DMT wire myographs (Danish MyoTech, Aarhus, Denmark). The MRA was oxygenated (95% $O_2$ and 5% $CO_2$), heated (37°C) and submerged in regular Krebs solution in mM: (NaCl 130; KCl 4.7; $NaHCO_3$ 14.9; $CaCl_2.2H_2O$ 1.56; $KH_2PO_4$ 1.18; $MgSO_4.7H_2O$ 1.18; EDTA 0.026; glucose 5.6, pH 7.40) to mimic an environment for optimal function. The MRA were normalized to their optimal lumen diameter for active tension development. To test vascular smooth muscle cell integrity, the arteries were initially contracted with modified 120 mM high potassium chloride (KCl) Krebs solution with the following composition in mM: (NaCl 14.7; KCl 120; $NaHCO_3$ 14.9; $CaCl_2.2H_2O$ 1.56; $KH_2PO_4$ 1.18; $MgSO_4.7H_2O$ 1.18; EDTA 0.026; glucose 5.6, pH 7.40). Cumulative concentration-effect curves to ACh (1 pM to 10 μM) after a previous contraction elicited by the thromboxane $A_2$ receptor analogue U46619 (30 nmol/L) was performed to evaluate vascular relaxation. Relaxation responses to ACh are shown as a percent of the initial U46619 contraction. Cumulative concentration-effect curve to the selective $\alpha_1$-adrenergic receptor agonist phenylephrine (PE; 0.1 nmol/L to 100 μmol/L) or to U46619 (1 pmol/L to 1 μmol/L) were also performed, and values were represented as milliNewton per millimeter (mN/mm).

**Plasma TNF-α**

Plasma TNF-α was measured using Mouse TNF-α high sensitivity ELISA kit (eBioscience, BMS607HS), according to the manufacturer's instruction. Plasma samples were obtained from male and female BPN/3J and BPH/2J LVCO- or LV-OSK-treated mice (i.v.). Microwell strips were washed twice with the manufacturer's wash buffer (400 μL each) and 50 μL of sample diluent was applied in all wells, and the external diluted standard (50 μL) was added in duplicate. After, 50 μL of each sample was added in duplicate and Calibrator Diluent was used as a blank condition, and biotin-conjugated (50 μL) was applied in all wells. The plate was sealed and incubated at room temperature for 2h in a microplate shaker. Subsequently, microwell strips were washed 6 times, and 100 μL diluted Streptavidin-HRP solution was added to all wells and incubated for 1 h at room temperature in a microplate shaker. After incubation, plates were washed 6 times, Amplification Solution I (100 μL) was added in all wells and incubated for 15 min at room temperature (shaking). Following, 6 washes were performed, Amplification Solution II was added to all wells and incubated at room temperature for 30 min in a microplate shaker. The plate was washed 6 times and TMB Substrate Solution was added (100 μL) to all wells for 20 min incubation at room temperature. Enzyme reaction was stopped by adding 100 μL Stop Solution and the plate was read on a spectrophotometer using 450 nm as the primary wavelength. Samples were obtained from 4 different animals processed at the same day.

**Plasma estradiol**

The estrogen levels in plasma samples were determined in female mice (40-44-week-old) i.v. treated with LVCO or LV-OSK using a commercial ELISA kit (Cayman Chemical, #501890), Briefly, the assay-specific reagents included estradiol acetylcholinesterase (AchE) tracer and estradiol ELISA antiserum. The 50 μL of samples and standard solutions were transferred to their respective wells. Subsequently, 50 μL of estradiol AchE tracer and 50 μL of estradiol antiserum were added to their designated wells, mixed gently, and then incubated for 60 min at room temperature. Afterward, the liquid was discarded, and the wells were washed four times with a washing buffer. Following the washing process, 200 μL of Elman's work reagent was added and incubated for 60 min at room temperature in the dark. At the end, the absorbance of the resulting reagent was measured at 405 nm, and data were presented as pg/mL. Samples were evaluated on duplicates from 5-6 different animals processed at the same day.

**Transmission electron microscopy (TEM)**

Cell and vascular morphologies were evaluated in mouse ECs or MRA. Briefly, mouse ECs treated with PBS, LVCO or LV-OSK for 3 days, or freshly dissected male mouse MRA i.v. treated with LVCO or LV-OSK (40-44 weeks of age) were mounted on wire myography for vascular reactivity assay, as aforementioned. After, MRA was carefully removed and fixed with 2.5% glutaraldehyde (pH 7.4) for at least 24 h. In addition, ECs were centrifuged at 300 xg at room temperature, and the pellet was fixed with 2.5% glutaraldehyde (pH 7.4) for at least 24 h. Subsequently, three washes in 0.1 M phosphate buffer followed by fixation with 1% osmium tetroxide and 1.5% potassium ferricyanide buffer for 1 h was performed on samples. After, specimens were dehydrated in ethanol, followed by acetonitrile incubations, embedded, and cured (48 h at 60° C). Ultrathin sections (80 nm) were cut on an ultramicrotome (Leica UltraCutR) and stained with 2% uranyl acetate and Hanaichi lead citrate, at room temperature. In addition, the morphological structure of LV-OSK was observed after staining with 2% uranyl acetate and Hanaichi lead citrate (negative staining), at room temperature, and compared to staining solutions only. Images were acquired with a JEOL 1400 Plus Transmission Electron Microscope.

**Multi-photon microscopy and second harmonic generation (SHG)**

MRA isolated from male LVCO- or LV-OSK- i.v. treated BPN/3J and BPH/2J mice (at 40-44 weeks of age) was investigated on their collagen, elastin content, and EGFP signals by second harmonic generation (SHG) signals using a multi-photon microscopy. Briefly, MRA was isolated from mice, cleaned off adjacent tissue, embedded in Tissue-Tek® O.C.T. Compound (Optimal Cutting Tissue; Sakura, #4583), frozen in dry ice, and stored at -80° C until experiments. Following, sample sections (5 μm) were incubated with PBS for 5 min at room temperature, and Fluoroshield™ with DAPI (Sigma Aldrich, F6057) was used as mounting media and for nuclei staining. Collagen, elastin, EGFP, and nuclei were visualized using Leica SP8 multiphoton MP system (Leica Microsystems, Mannheim, Germany), with pulsed femtosecond Titanium:Sapphire (Ti:Sa) laser (Chameleon Vision II, Coherent, Santa Clara, CA, USA) tunable for 880 nm (for collagen), and 720 nm (for DAPI). Multi-photon excitation at 880 nm was used to reveal structural information of collagen fibers by intrinsic contrast imaging of the SHG signal. Elastin autofluorescence and EGFP signals were observed at 488 nm (498 to 550 nm). Images were acquired on a 25x water objective, using xyz series on a sequential mode with high resolution (1024x1024 format, 400 Hz, 16 bits), and 3D reconstructions were automatically performed with LAS X Software. Images were analyzed by FIJI Software using max projections of z series, and percentage (%) of collagen area, or elastin+EGFP$^+$ area were analyzed. The total area corresponded to 442.86 μm$^2$ at 25x objective. Negative control represented photos obtained from MRA isolated from male BPN/3J mice with no treatments.

### Statistical analysis

All statistical analyses were performed using GraphPad Prism 9.0 (GraphPad Software Inc., La Jolla, CA, USA). Data are presented as mean ± standard error of the mean (S.E.M) and statistical significance was set at $p<0.05$. All bar graphs are accompanied by dispersion of the individual values (n). The pharmacological parameters of agonist potency ($pD_2$; negative logarithm of agonist $EC_{50}$) and maximum effect (ME, in %) were calculated for each agonist using the non-linear curve regression of the specific cumulative concentration-effect curves. The unpaired and two-tailed Student's t test was used to compare two different groups, or the One-way or two-way analysis of variance (ANOVA), followed by Tukey post hoc was used to identify interaction factors and compare 3 or more groups. The sample size (n) indicated per experiment is the number of independent samples used.

### Supplemental Figures and Figure Legends

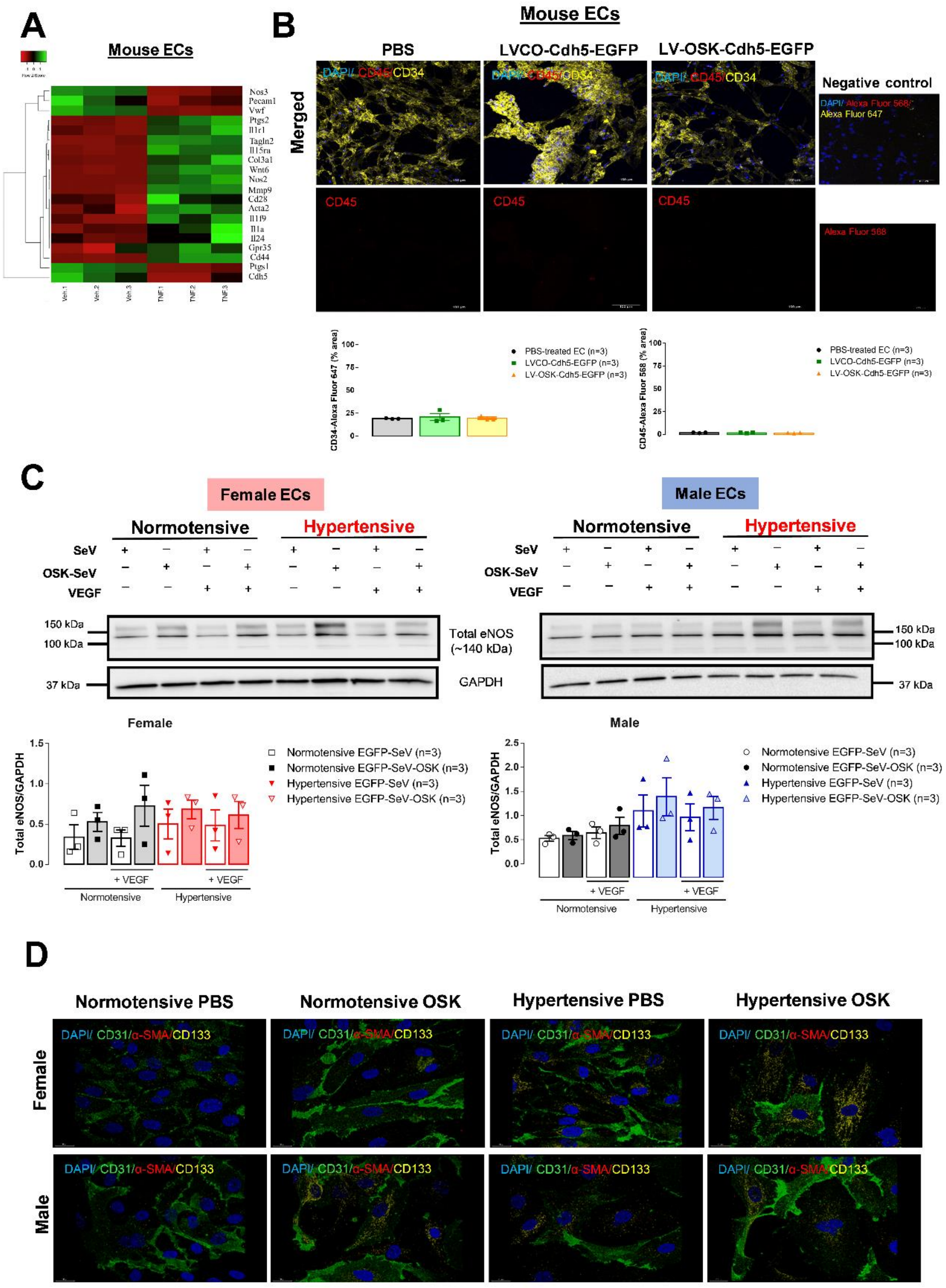


**Figure S1**: **TNF-α induced EndMT genes in mouse ECs**. In **A**, heatmap of main EndMT genes in mouse ECs treated with PBS (vehicle) or TNF-α (100 nM, 24 h; n=3).

In **B**, representative images, and bar graphs show that CD34 staining (yellow) was not different between PBS, LVCO or LV-OSK treatment in mouse ECs (upper panel; merged images), and CD45 (red) was not detected (lower panel) ($p>0.05$, n=3). DAPI (blue) represented nuclei staining. Negative controls were cells with no primary antibody incubation and DAPI, and bar represents 50 μm. In **C**, the OSK-SeV treatment did not change total eNOS protein expression (~140 kDa) in female (left) or male (right) HAoECs ($p>0.05$, n=3), in basal condition or after VEGF (2 ng/mL, 2 h). GAPDH was used as loading control. In **D**, representative 3D reconstruction images (merged) show DAPI (blue), CD31 (green), α-SMA (red) and CD133 (yellow) in human ECs from normotensive and hypertensive patients, treated with PSB or OSK-SeV (n=1). Bar represents 20 μm (63x objective).

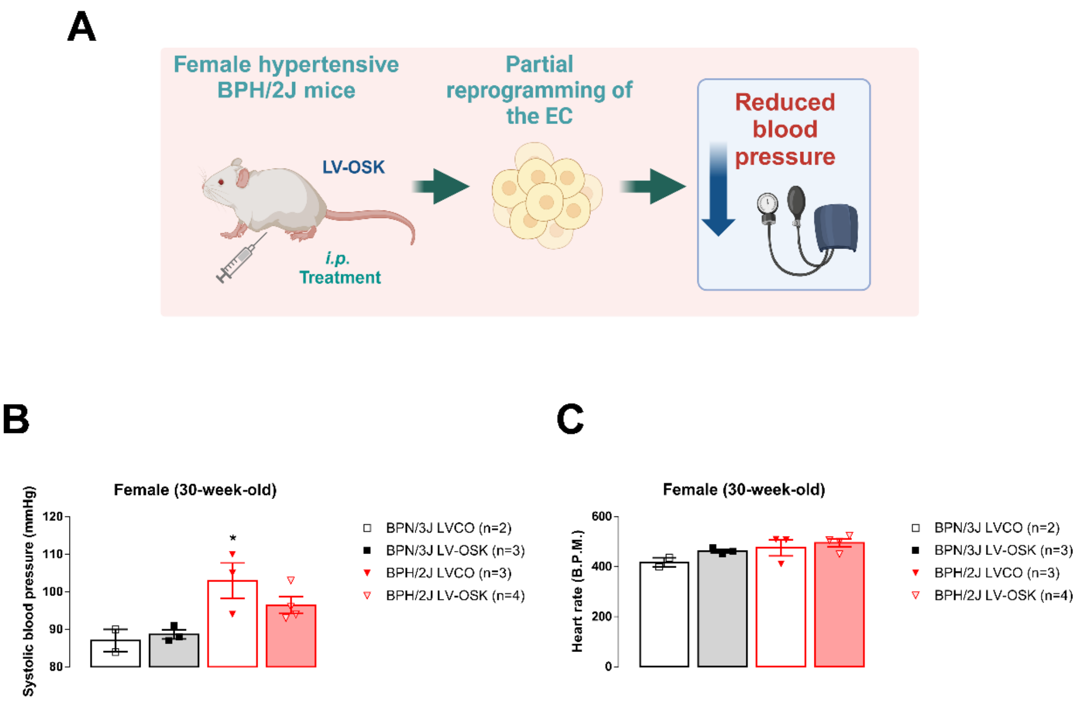


**Figure S2**: **Early treatment of female BPH/2J with OSK tended to decrease blood pressure**. In **A**, graphical abstract for the experimental design of *in vivo* intraperitoneal (i.p.) treatment of female BPN/3J or BPH/2J mice at 30 weeks of age (created with BioRender.com). In **B** and **C**, systolic blood pressure (mmHg) and heart rate (B.P.M.) values measured 10 days after i.p. treatment with LVCO or LV-OSK in female BPN/3J and BPH/2J mice at 30 weeks of age (p<0.05; * different from BPN/3J LVCO mice, n=2-4).

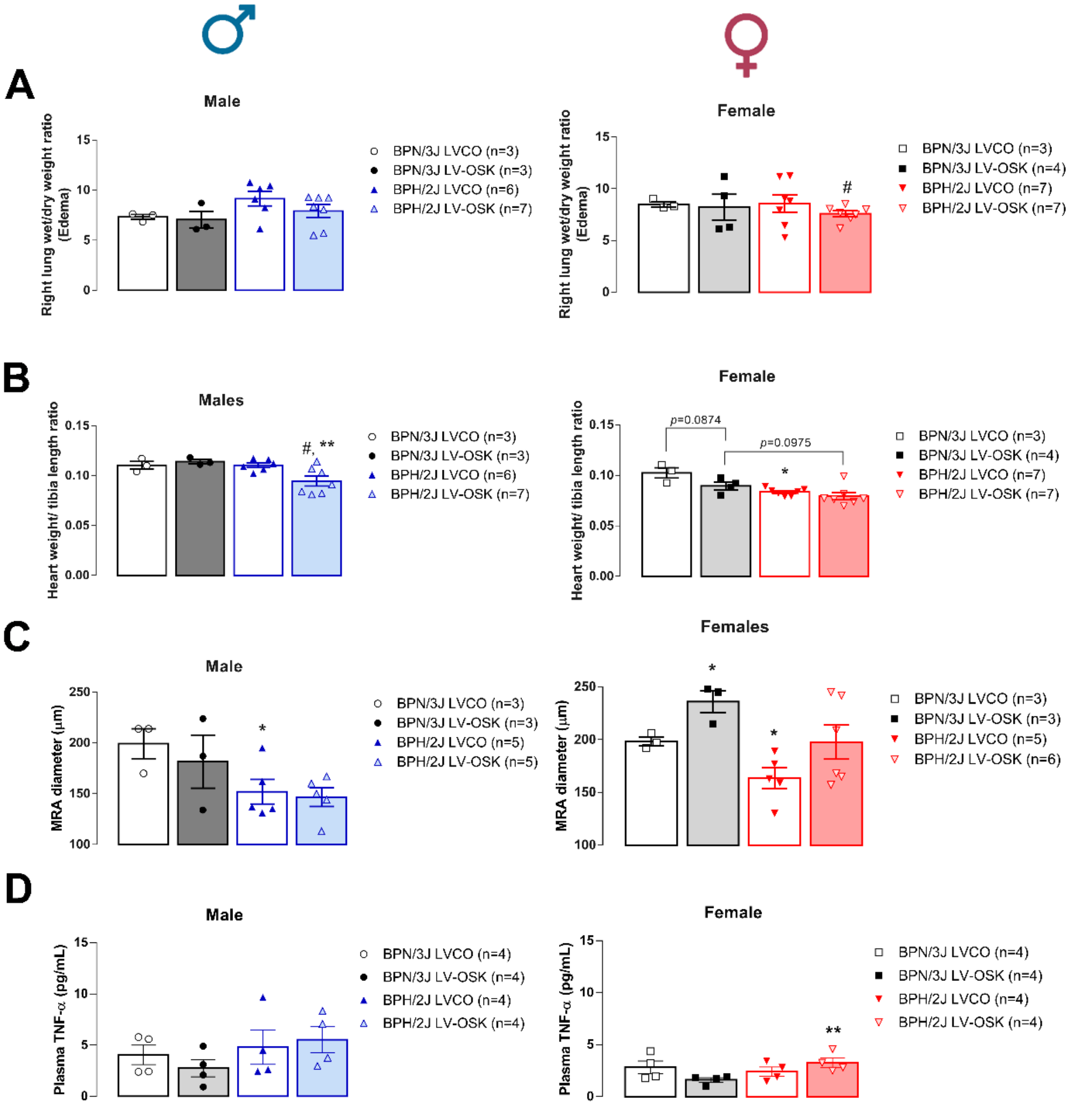


**Figure S3**: **EC *in vivo* reprogramming did not induce lung edema in mice**. In **A**, bar graphs show that the i.v. LV-OSK treatment did not induce lung edema in male (left) or female (right) mice at 40-44 weeks of age. Heart weight/tibia length ratio (**B**) was lower in male, but not in female BPH/2J mice treated with LV-OSK compared to other groups ($p<0.05$, n=3-7). MRA inner diameter (**C**) was lower in male and female LVCO-treated BPH/2J compared to LVCO-treated BPN/3J mice. MRA diameter in female LV-OSK-treated BPN/3J was higher compared to female LVCO-treated BPN/3J mice. Plasma TNF-α levels (**D**) did not modify in male but increased in female BPH/2J mice treated with LV-OSK ($p<0.05$; * different from BPN/3J LVCO mice; # different from BPH/2J LVCO mice; ** different from BPN/3J LV-OSK mice, n=3-7).

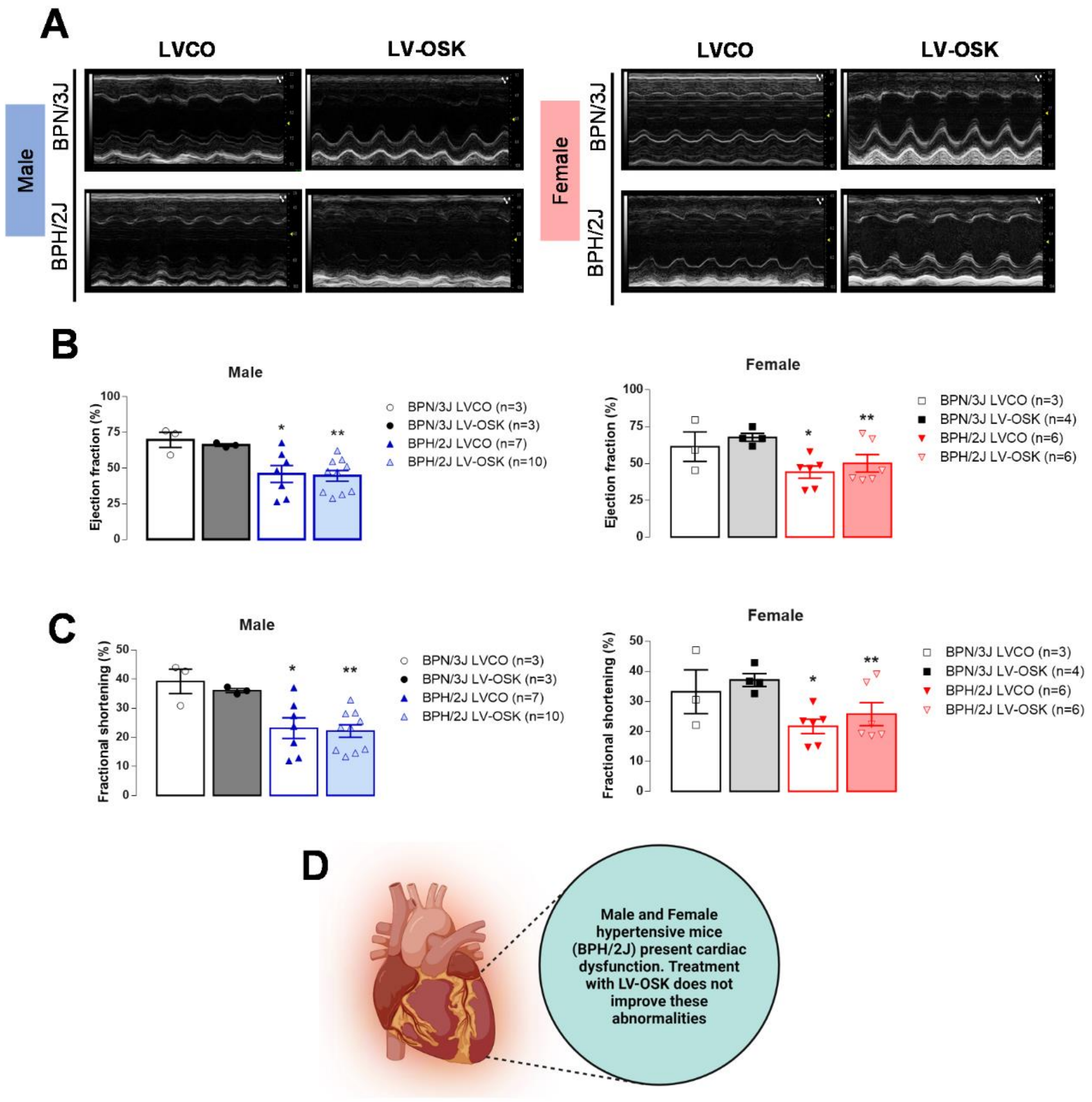


**Figure S4**: Cardiac function was not ameliorated by LV-OSK treatment in mice. Representative M-mode short axis images (**A**) obtained from LVCO- or LV-OSK-treated male and female BPN/3J and BPH/2J mice at 40-44 weeks of age. While ejection fraction and fractional shortening were not different in male and female BPN/3J mice, these parameters were reduced in BPH/2J mice (**B** and **C**) regardless of sexes. LV-OSK treatment did not improve cardiac function in neither male nor female mice compared to their respective LVCO-treated BPH/2J mice. In **D**, graphical abstract summarizes the cardiac dysfunction in male and female hypertensive mice, and the lack of cardiac effects induced by LV-OSK treatment (created with BioRender.com) ($p<0.05$; * different from BPN/3J LVCO mice; ** different from BPN/3J LV-OSK mice, n=3-10).

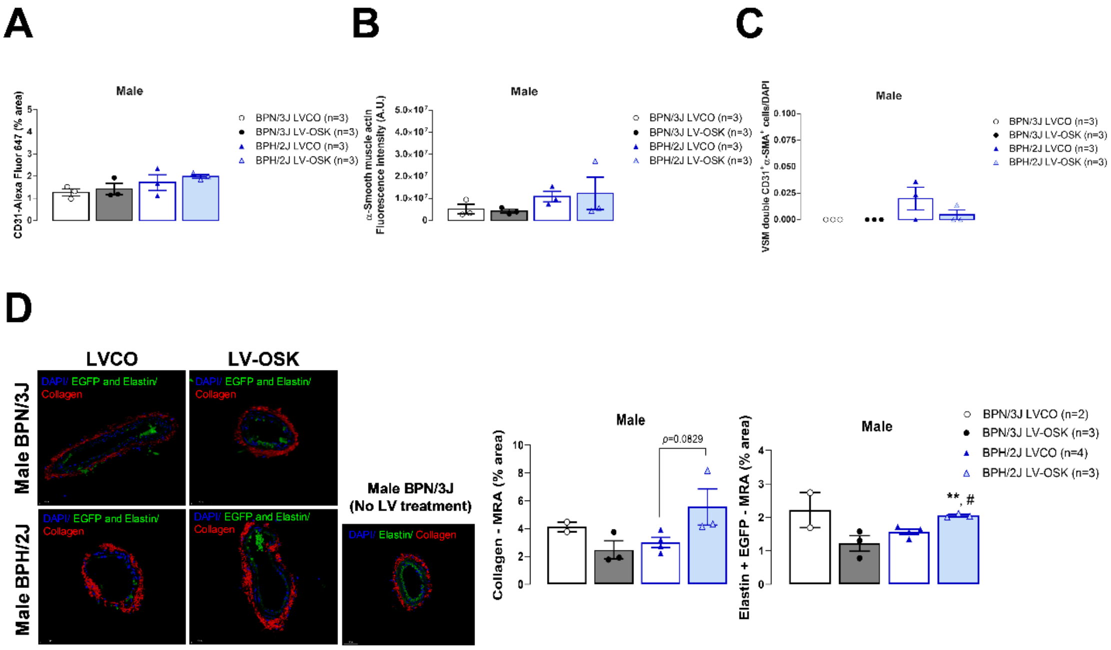


**Figure S5**: **EC *in vivo* reprogramming did not change CD31 or α-SMA expression in male BPH/2J mice aortas**. The i.v. LV-OSK treatment did not change the total vascular endothelium area of CD31 (**A**; %) or α-SMA (**B**; A.U.) in male aortas at 40-44 weeks of age. In **C**, bar graph shows that the double positive $CD31^+$ and $\alpha$-$SMA^+$ cells/DAPI ratio in the vascular smooth muscle (VSM) layer from mouse aortas were not different between the groups ($p>0.05$, n=3). In **D**, 3D reconstruction of SHG images for MRA from male BPN/3J and BPH/2J mice show nuclei (DAPI, blue), collagen (red), elastin and $EGFP^+$ cells (green). Images were acquired on multi-photon SP8 microscope (TCS SP8 MP Leica), and the negative control represents MRA from male

BPN/3J mice with no treatments and incubated with DAPI. Bar represents 50 µm. Bar graphs show collagen (%) and elastin and EGFP+ cells (%) quantification in MRA, respectively ($p<0.05$; # different from BPH/2J LVCO; ** different from BPN/3J LV-OSK, n=2-4).

**Supplemental Table S1**. **Cardiac function measured by echocardiography in male or female BPN/3J and BPH/2J mice at 40-44 weeks of age, after i.v. LVCO or LV-OSK treatment**. LV-OSK did not ameliorate the cardiac function or increased tibia length values in male or female BPH/2J mice.

| Parameters | Male BPN/3 | Male BPN/3J | Male BPH/2J | Male BPH/2J | Female BPN/3J | Female BPN/3J | Female BPH/2J | Female BPH/2J |
|---|---|---|---|---|---|---|---|---|
| | LVCO | LV-OSK | LVCO | LV-OSK | LVCO | LV-OSK | LVCO | LV-OSK |
| Diameter systole (mm) | 2.26 ± 0.21, n=3 | 2.50 ± 0.03, n=3 | 2.98 ± 0.21, n=7 | 3.09 ± 0.17, n=10 | 2.26 ± 0.34, n=3 | 2.30 ± 0.12, n=4 | 2.98 ± 0.13, n=6 | 2.99 ± 0.20, n=6 |
| Diameter diastole (mm) | 3.70 ± 0.14, n=3 | 3.91 ± 0.01, n=3 | 3.86 ± 0.16, n=7 | 3.95 ± 0.14, n=10 | 3.35 ± 0.16, n=3 | 3.66 ± 0.14, n=4 | 3.80 ± 0.08, n=6 | 4.01 ± 0.08, n=6 |
| Volume systole (uL) | 17.89 ± 4.05, n=3 | 22.42 ± 0.68, n=3 | 6.24 ± 6.38, n=7 | 39.29 ± 5.06, n=10 | 18.82 ± 6.58, n=3 | 18.40 ± 2.43, n=4 | 34.99 ± 3.58, n=6 | 35.99 ± 5.35, n=6 |
| Volume diastole (uL) | 58.56 ± 5.31, n=3 | 66.33 ± 0.24, n=3 | 65.42 ± 6.85, n=7 | 68.99 ± 5.70, n=10 | 46.30 ± 5.11, n=3 | 56.80 ± 5.42, n=4 | 62.25 ± 3.22, n=6 | 70.51 ± 3.41, n=6 |
| Stroke volume (uL) | 40.66 ± 3.95, n=3 | 43.92 ± 0.46, n=3 | 29.17 ± 3.58, n=7 | 29.70 ± 2.63, n=10** | 27.49 ± 1.77, n=3 | 38.40 ± 3.69, n=4 | 27.27 ± 2.55, n=6 | 34.52 ± 2.62, n=6 |
| Cardiac output (mL/min) | 15.97 ± 1.51, n=3 | 17.81 ± 1.18, n=3 | 12.18 ± 1.55, n=7 | 12.52 ± 0.91, n=10 | 11.05 ± 1.56, n=3 | 16.03 ± 2.13, n=4 | 11.36 ± 1.27, n=6 | 13.99 ± 0.91, n=6 |
| Left ventricle mass (mg) | 245.67 ± 36.0, n=3 | 278.34 ± 34.72, n=3 | 183.38 ± 20.94, n=7 | 168.29 ± 8.22, n=10** | 203.25 ± 11.86, n=3 | 162.29 ± 6.41, n=4 | 117.28 ± 8.17, n=6* | 134.95 ± 8.96, n=6 |
| LVAW systole (mm) | 1.97 ± 0.17, n=3 | 1.76 ± 0.08, n=3 | 1.44 ± 0.13, n=7* | 1.34 ± 0.07, n=10 | 1.53 ± 0.17, n=3 | 1.58 ± 0.04, n=4 | 1.23 ± 0.06, n=6 | 1.28 ± 0.08, n=6 |
| LVAW diastole (mm) | 1.49 ± 0.17, n=3 | 1.26 ± 0.10 n=3 | 1.12 ± 0.11, n=7 | 0.98 ± 0.07, n=10 | 1.26 ± 0.19, n=3 | 1.12 ± 0.06, n=4 | 0.89 ± 0.05, n=6* | 0.91 ± 0.07, n=6 |
| LVPW systole (mm) | 1.79 ± 0.24, n=3 | 2.11 ± 0.42, n=3 | 1.40 ± 0.12, n=7 | 1.36 ± 0.07, n=10** | 1.93 ± 0.11, n=3 | 1.51 ± 0.04, n=4 | 1.03 ± 0.06, n=6* | 1.18 ± 0.12, n=6 |
| LVPW diastole (mm) | 1.36 ± 0.20, n=3 | 1.69 ± 0.30, n=3 | 1.15 ± 0.12, n=7 | 1.13 ± 0.06, n=10** | 1.52 ± 0.07, n=3 | 1.10 ± 0.05, n=4* | 0.80 ± 0.04, n=6* | 0.85 ± 0.06, n=6** |
| LVID systole (mm) | 2.17 ± 0.20, n=3 | 2.41 ± 0.04, n=3 | 2.79 ± 0.21, n=3 | 3.38 ± 0.29, n=4 | 2.19 ± 0.36, n=3 | 2.49 ± 0.08, n=3 | 3.18 ± 0.19, n=5* | 3.32 ± 0.022, n=3 |
| LVID diastole (mm) | 3.56 ± 0.16, n=3 | 3.90 ± 0.02, n=3 | 3.66 ± 0.04, n=3 | 4.11 ± 0.20, n=4 | 3.30 ± 0.16, n=3 | 3.67 ± 0.13, n=3 | 3.85 ± 0.15, n=5 | 4.12 ± 0.10, n=3 |
| PWV (mm/ms) | 1.09 ± 0.12, n=3 | 1.29 ± 0.23, n=3 | 2.39 ±0.79, n=7 | 2.39 ± 0.83, n=7 | 1.44 ± 0.30, n=3 | 1.01 ± 0.13, n=4 | 1.66 ± 0.34, n=6 | 3,83 ± 2.26, n=6 |
| Tibia length (cm) | 1.90 ± 0.06, n=3 | 1.83 ± 0.03, n=3 | 1.70 ± 0.03, n=6* | 1.69 ± 0.03, n=7** | 1.97 ± 0.03, n=3 | 1.95 ±0.07, n=4 | 1.70 ± 0.03, n=7* | 1.83 ± 0.07, n=7 |

* Different from male or female BPN/3J LVCO; ** different from male or female BPH/2J LVCO; Two-way ANOVA, followed by Tukey post-hoc, $p<0.05$; n=3-10. LVAW means left ventricle anterior wall; LVPW means left ventricle posterior wall; LVID means left ventricle inner dimension; and PWV means pulse wave velocity calculated in the aortic arch.
