## Supplementary material for "A Single-Short Partial Reprogramming of the Endothelial Cells decreases Blood Pressure via attenuation of EndMT in Hypertensive Mice": Unedited WB membranes

### Uncropped gels

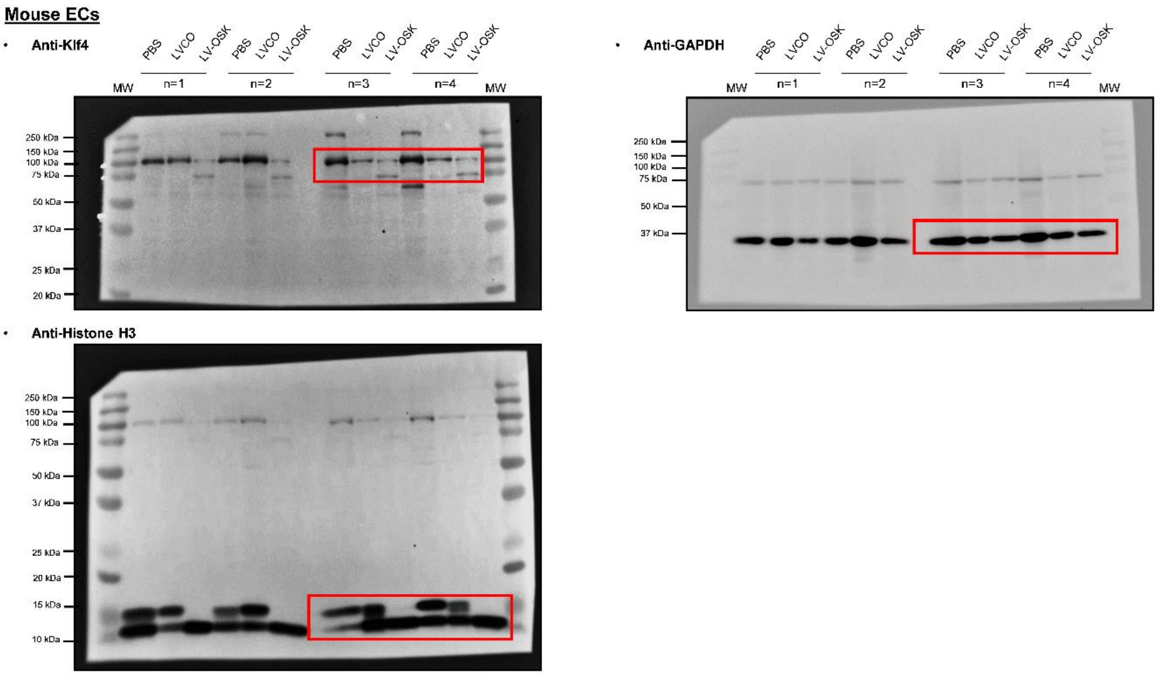

Female HAoEC: eNOS, p-eNOS (Ser1177) and GAPDH

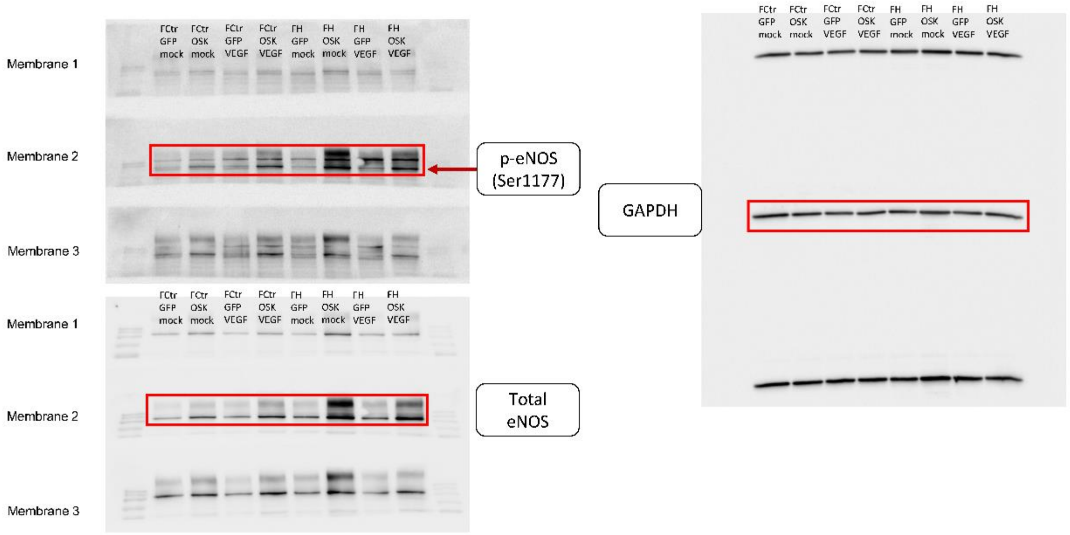


Male HAoEC: eNOS, p-eNOS (Ser1177) and GAPDH

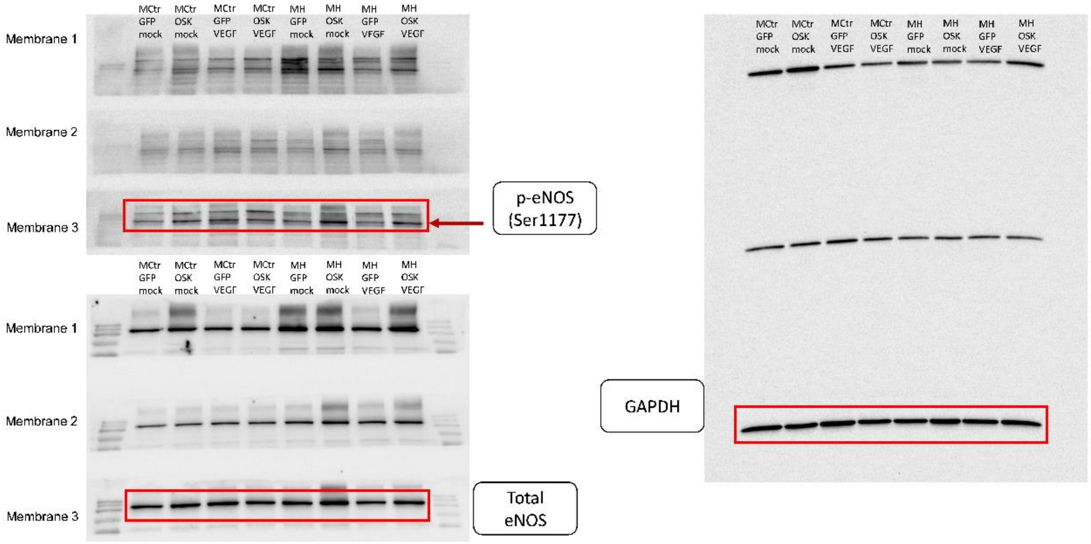
